# Angiotensin (1-7)-based Proteomimetic Polymers as Topicals for the Remediation and Repair of the Cornea following Injury

**DOI:** 10.64898/2026.09.30.755738

**Authors:** Parisa Foroozandeh, Ashley K. Nensel, Xiaolin Qi, Jessica Marks, Nihal Kaplan, Siwon Park, Xiaoping Chen, Amit Fischer, Maria Telesforo, Robert M. Lavker, Nathan C. Gianneschi, Han Peng

**Author notes:** Corresponding authors: Prof Robert Lavker; Prof Han Peng; Prof Nathan Gianneschi. Denotes equal author contributions (Parisa Foroozandeh, Ashley K. Nensel).

## Abstract

The cornea is the outermost layer of the eye, most vulnerable to injury including from chemical exposure, either environmental, industrial or chemical weapons. For example, exposure of the eye to sulfur mustard (SM) can cause mustard gas keratopathy, resulting in devastating consequences for vision. Available treatments for corneal mustard keratopathy are either ineffective, or if used for a prolonged time, have deleterious side effects. Here, we aimed to harness the anti-inflammatory and anti-angiogenic peptide angiotensin-(1-7), known as Ang-(1-7), that is derived from the renin-angiotensin system (RAS). We designed and developed a protein-like polymer (PLP) displaying a brush of norleucine modified Ang (1-7) peptides (NLE Ang (1-7) PLP), which is a more potent derivative of Ang-(1-7), for administration as an eye drop. In a mouse model of nitrogen mustard (NM)-induced corneal injury, topical treatment and intraperitoneal (i.p.) injection of NLE Ang (1-7) PLP persistently minimized: (i) corneal haze; (ii) edema; (iii), expression of pro-inflammatory and pro-angiogenic genes; and (iv) neovascularization and conjunctivalization as evidenced after a prolonged treatment and recovery period. Collectively, our observations have demonstrated the efficacy of this novel polymer proteomimetic based eye drop treatment in reducing chemical agent-induced corneal injury as well as indicating its potential application in other ocular surface diseases associated with inflammation and angiogenesis.

## Introduction

The anterior surface of the eye functions as a barrier to the external environment and protects the delicate underlying tissues from injury. Central to this protection are the cornea, limbus and conjunctiva, in conjunction with the tear film. The corneal epithelium, which is self-renewing stratified squamous epithelium, supports a tear film and helps maintain transparency so that light can be transmitted to the interior of the eye^1–8^ . One of the features of the cornea is that following a relatively superficial debridement wound, the stromal surface is covered with a provisional epithelium within a day. This process is regulated via limbal epithelial stem cells (LESCs) preferentially located as a subpopulation of limbal basal epithelial cells^2,9^. Superficial corneal epithelial wounds are not associated with a significant inflammatory response. However, if the wound is more substantial, with a breach of the basement membrane, a brisk stromal inflammation occurs. Immune cell recruitment after corneal injury is mediated by proinflammatory cytokines released from epithelial cells and keratocytes at the injured site. For example, interleukin-1(IL-1), interleukin-6 (lL-6) and tumor necrosis factor receptor (TNF) ^10,11^ aid in attracting neutrophils, which are the first cells infiltrating the cornea after injury^12,13^. Shortly after neutrophils enter the cornea, macrophages extravasate from the limbal vessels, infiltrate the stroma from superficial to deeper layers and migrate towards the center of the cornea. In addition to removal of debris and apoptotic cells, macrophages are essential mediators of angiogenesis after severe and prolonged corneal injury^14^. Several chemokines and their receptors have been identified in the inflamed cornea. CXCL1, CXCL2, CXCL5, CXCL8, and CCL2 mRNA levels were found to be elevated in human and/or mouse inflamed corneas^15–18^. Finally, the corneal angiogenic privilege is essential for good vision and is therefore strictly regulated. This condition can be over-ridden following a severe inflammatory event resulting in upregulation of proangiogenic growth factors^18^. This leads to a secondary ingrowth of both blood and lymphatic vessels from the limbal area into the central cornea. Such neovascularization adversely affects the transparency of the cornea and compromises vision. Consequently, after the immune response is complete and barrier function has been achieved, corneal blood and lymphatic vessels need to resolve promptly to restore corneal transparency. However, in certain pathological conditions (e.g., sulfur mustard injury) corneal blood and lymphatic vessel regress slowly, and may persist thus contributing to compromised vision ^19–24^.

Sulfur mustard (SM), the “king of chemical warfare agents,” is a blistering compound that can alkylate DNA and RNA strands, breakdown proteins and lipids of the cell membrane resulting in cell death^24^. The eyes are arguably one of the most important organs that can be damaged by contact with sulfur mustard. The degree of ocular damage varies, and it is estimated that alterations are noted in 50-90% of individuals exposed to SM^24^. Even small amounts of SM can cause visual impairment and disability^25–28^. Following exposure, there is an acute and a delayed phase. In both humans and animals, the acute stage of injury lasts 1 to 2 weeks and includes conjunctivitis, photophobia, and corneal swelling^27,29–31^.

In both humans and animal models exposed to SM, following a period of a few weeks to months of clinical quiescence, chronic or delayed injury can arise. This is characterized clinically as corneal haze, extensive edema, vascularity, and corneal epithelial defect. The degree of corneal epithelial defect is more extensive than seen during acute injury. The profoundly disorganized stroma is accompanied by scaring and/or hazing. A significant increase in cytokines (IL1β, TNFα, IL6, IL8), chemokines and MMPs (MMP2 and 9) are a feature of the injury^32^. Neovascularization often accompanies keratitis and harbors an unfavorable outcome for vision. Finally, it is recognized that limbal stem cell deficiency (LSCD) may derive directly from the destructive loss of LESC or indirectly from an alteration in the stem cell niche due to the persistent inflammation^29,33^. Therefore, countermeasures for corneal mustard exposure should have the ability to halt the loss of epithelial cells particularly LESCs.

With respect to treatment modalities, either steroidal or non-steroidal anti-inflammatory drugs are accepted treatments for the acute and prolonged phases. While effective in combatting inflammation, steroids are contra-indicated for epithelial wound healing^34^. Specifically, there have been warnings against the use of topical steroids in SM ocular injuries^35^. Furthermore, it is well-recognized that the many adverse side effects of long-term steroid use limit their effectiveness^34^. For those victims who have bacterial infections resulting from SM injury, antibiotics such as sulfacetamide, gentamycin, or neomycin have been used. However, these antibiotics have limited efficacy in treating corneal erosions. Thus, it is necessary to develop novel topical treatments for mustard keratopathy that act during the acute and delayed phases to prevent ocular deterioration and facilitate resolution of corneal inflammation.

Corneal wound healing and inflammation have previously been shown to be mediated by the renin-angiotensin system (RAS) in mouse models^36–39^. RAS relies on two axes that exert both traditional pro-inflammatory effects and an anti-inflammatory counterbalance cascade^40^. The traditional pro-inflammatory axis uses the Angiotensin-converting enzyme (ACE) to cleave Angiotensinogen (1-225) to the active Angiotensin II (1-8) peptide which binds the Angiotensin II Type I Receptor (ATR1). However, to exert regulatory anti-inflammatory and anti-proliferative effects, a second enzyme ACE2 will cleave one additional residue to form anti-angiogenic peptide angiotensin-(1-7) (Ang-(1-7)) which binds a separate MAS receptor and stimulates the development of progenitor cells and vasodilation^41^. ACE2 has an anti-inflammatory role, due to its ability to catalyze and produce Ang-(1-7) in a variety of tissues including the cornea^37^. Studies have shown that exogenous delivery of Ang (1-7) peptide accelerates dermal wound healing by modulating cell proliferation and angiogenesis^36,42–47^. A more potent analogue of this peptide, NLE Ang (1-7), has accelerated healing properties, reduction of fibrotic scars and normalization of tissue post mechanical injury in diabetic skin and cornea^36,44–48^ . However, peptides suffer from several key disadvantages. Topical delivery of peptides is difficult due to both their proteolytic instability and their tendency to aggregate which prevents penetration across biological membranes.

Herein, we have harnessed the anti-inflammatory properties of the NLE Ang (1-7) peptide while enhancing therapeutic half-life and multivalency through their incorporation into a Protein-like Polymer (PLP), which is a proteomimetic bottlebrush polymer ^49^. PLPs are synthesized by modifying peptides to include a polymerizable norbornene at either the N- or C-termini. The resulting monomers are then subjected to graft-through ring opening metathesis polymerization (ROMP) ^50^ to prepare brush polymers of functional peptides ^51–53^. This process yields low dispersity polymers with tunable molecular weight and facile, modular incorporation of functional handles (i.e. fluorescent dyes, biotin). Previous reports have shown that these polymers have extended circulation half-lives *in vivo*, exhibit proteolytic stability, and show multivalent engagement to protein and cellular targets ^54–58^. By harnessing these properties, we demonstrate that NLE Ang (1-7) PLP accelerates healing from chemical injury in the eye. Previous work using anti-angiogenic PLPs, derived from Thrombospondin-1, have been applied successfully to the eye in models of neovascular age-related macular degeneration (nAMD) but required intravitreal injections for efficacy ^58^. Here, we formulated PLPs as eye drops, because they are clearly less invasive for the patient and appropriate for this injury model. Therefore, we compare the chemical injury wound healing efficacy of the peptide analogue and the NLE Ang (1-7) PLP administered through both systemic (intraperitoneal) and topical delivery (eye drop). Our findings demonstrate that this novel NLE Ang (1-7) PLP based eye drop has significant efficacy in treating mustard agent-induced corneal injury as well as highlights its broader potential for treating other ocular surface diseases associated with inflammation and angiogenesis.

## Materials and Methods

All animal experiments were performed at Northwestern University Feinberg School of Medicine using only protocols approved by the Institutional Animal Care and Use Committee (IACUC) and complied with the ARVO Statement for the Use of Animals in Ophthalmic and Vision Research. Specific pathogen-free mice were purchased from commercial suppliers approved by the Center for Comparative Medicine at Northwestern University (CCM). Detailed descriptions of the PLP synthesis, characterization, enzyme stability, cell cultures, confocal microscopy, NM-injury models, histological staining, RT-qPCR, and ICP-MS are provided in *SI Appendix*.

## Results

### Synthesis and characterization of Ang(1-7) inspired PLPs and control PLPs

The peptide NLE Ang (1-7) was synthesized using standard solid phase peptide synthesis ^59,60^. This derivative of Ang (1-7), sequence DRVYIHP, replaces the valine in the third position to the unnatural amino acid norleucine (NorLeu, NLE), sequence DR(NorLeu)YIHP (Fig. 1, Fig. S1, Fig. S2, Table S1). Monomers were polymerized via graft through ROMP to produce PLPs with a target degree of polymerization of 15 (Fig. S3). After the conversion of monomers, additional functional groups were incorporated as either additional monomers (NorCy5) or terminating agents (Gd-Ta). Polymers were characterized using size-exclusion chromatography with multi-angle light scattering (SEC-MALS) to determine the molecular weight (Mn), degree of polymerization (DP), and dispersity (Đ). Molecular weight was also confirmed using SDS-PAGE and MALDI (Fig. S4). Dynamic light scattering showed dispersed 4 nm diameter PLPs (1 mg/ml in saline, Fig. S4C).

**Figure 1.**
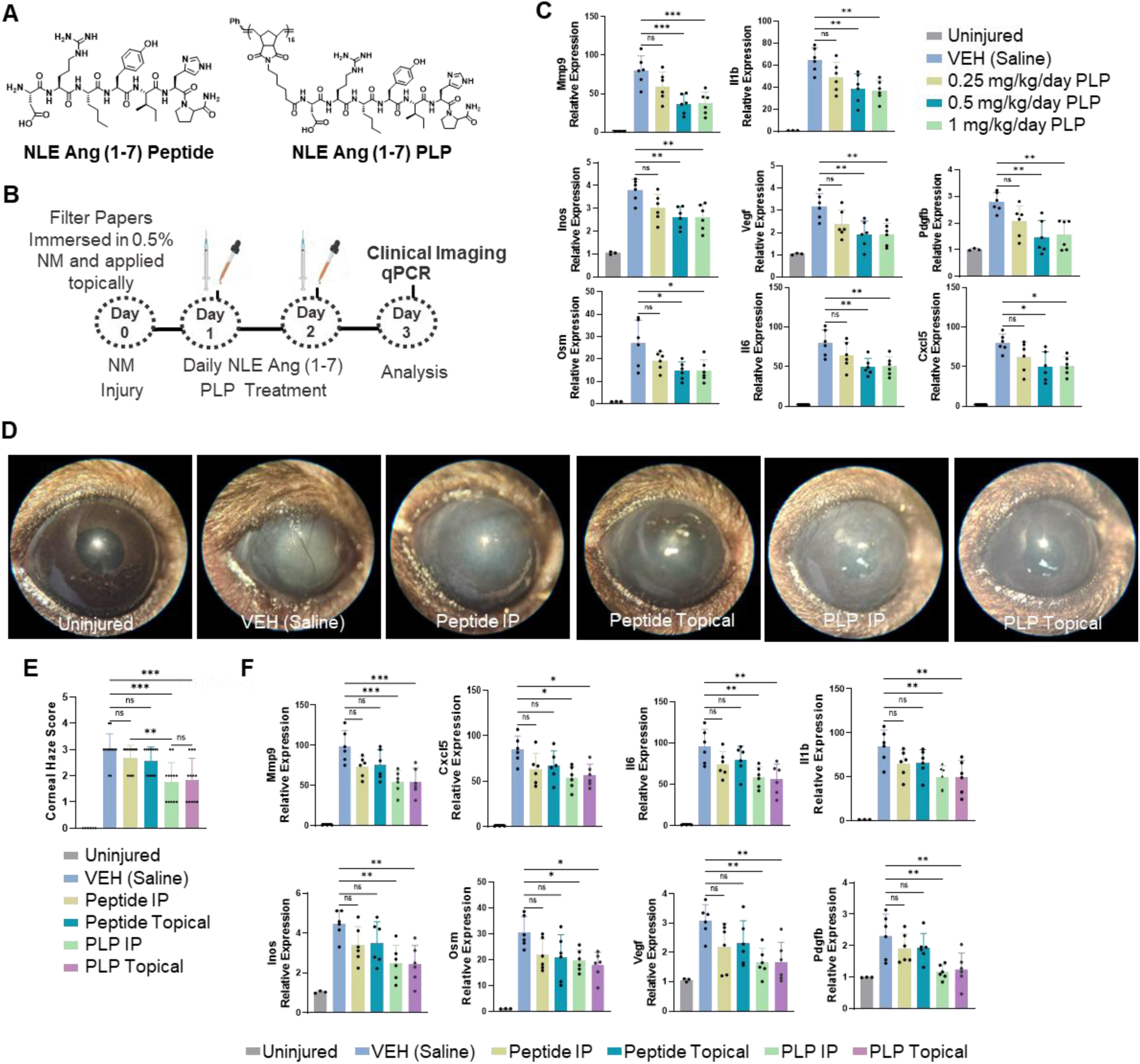
NLE Ang (1-7) PLP treatment attenuates corneal haze and inflammation in acute phase of NM injury. (A) Synthetic library of NLE Ang (1-7) compounds including the peptide of interest and its polymerized form. (B) Schematic representation of 3 day treatment. Mice eyes were exposed to 0.5% nitrogen mustard (NM) for one minute. Twenty-four hours post NM injury, mice were i.p. injected or topically treated with NLE Ang(1-7) PLP or NLE Ang(1-7) peptide once daily for 2 days (acute phase treatment, n=6 mice). (C) Dose-response (i.p. injection) of the NLE Ang(1-7) PLP was first assessed via the reduction of pro-inflammatory genes using RT-qPCR. (D-E) Routes of administration of the NLE Ang(1-7) PLP were next assessed. Corneal haze was (D) imaged and (E) scored at the end of the experiment. Topical treatment and i.p. injection of NLE Ang(1-7) PLP markedly decreased NM-induced corneal haze during acute phase of NM injury compared to vehicle. (F) The corneas were harvested, and RT-qPCR was conducted to determine the expression of pro-inflammatory genes as well as genes related to angiogenesis in mouse cornea (n=6). *: P ≤ 0.05, **:P ≤ 0.01, ***P ≤ 0.001, **** P ≤ 0.0001.

### Ang(1-7)-based PLPs show extended enzyme stability

Enzyme stability of NLE Ang (1-7) PLP and peptide were assessed (Fig. S5). Both compounds were dissolved at the same concentration with respect to peptide (100 μM) and treated with trypsin. Aliquots of the trypsin-treated compounds were taken, quenched, and analyzed using HPLC. A cleaved peptide fragment peak appears and increases across the week for both peptide and PLP. The area under the curve is measured and quantified via standard curve and correlated to the sample degradation (Fig. S6). Using this HPLC technique, peptide degradation after 24 hours indicated that 65% remained intact compared to 90% of the NLE Ang (1-7) PLP. Following a week of enzymatic incubation, this difference further increases with approximately 50% versus 85% of peptide to NLE Ang (1-7) PLP intact (Fig. S6C). This demonstrates that NLE Ang (1-7) PLP is significantly more stable than peptides.

### NLE Ang (1-7) PLPs penetrate corneal cells

We investigated whether NLE Ang (1-7) PLP can be internalized by corneal epithelial cells. Uptake studies were performed using fluorescently labeled PLPs. Here, hTCEpi cells, a limbal derived corneal epithelial cell line, were incubated with NorCy5-labeled NLE Ang (1-7) PLP (5, 1, or 0.5 µM) for 24 hours. Confocal microscopy was performed and fluorescent puncta were detected in cells, indicating that this corneal epithelial cell line can uptake PLP *in vitro* (Fig. S7).

Following nitrogen mustard (NM) injury, NorCy5-labeled NLE Ang (1-7) PLP was administered via eye drops and penetration across the cornea was assessed at time points of 1 hr., 4 hr., 8 hr., and 24 hr. Here, PLP-based fluorescence was detected only at the shortest time point of 1 hr. across the surface of the cornea (Fig. S8).

### Intra-peritoneal injection of NLE Ang (1-7) PLPs attenuate NM-induced corneal inflammation during the acute phase

We tested the efficacy of different doses of NLE Ang (1-7) PLP in resolving inflammation during the acute phase of NM-induced corneal injury (Fig. 1). One day after topical NM exposure to the cornea, mice were injected once daily with NLE Ang (1-7) PLP (Fig. 1A) for three days via intraperitoneal (i.p.) injection (Fig. 1B). Three days after NM exposure, RT-qPCR showed an increased expression of genes involved in inflammation (e.g., Mmp9, Cxcl5, IL1b, IL-6, Osm) and regulation of angiogenesis (e.g., Vegf, Pdgfb) (Fig. 1C, Table S2). Such increases in the expression of pro-inflammatory and pro-angiogenic genes were reversed with various doses of NLE Ang (1-7) PLP. Among the different doses used, treatment with NLE Ang (1-7) PLP at 0.5 mg/kg/day for two days showed the most pronounced reduction in pro-inflammatory and pro-angiogenic genes compared with both 1 mg/kg/day and 0.25 mg/kg/day doses.

### Topical treatment of NLE Ang (1-7) PLPs attenuate NM-induced corneal inflammation during acute phase

Since topical treated NLE Ang (1-7) PLPs can be internalized by corneal cells, we developed an eyedrop based on NLE Ang (1-7) PLPs and tested the efficacy of this eyedrop in corneal inflammation. One day after NM exposure, mice were topically treated with this eyedrop once daily for two days. To compare against i.p. injection, the same dose (0.5 mg/kg/day) was used for both i.p. injection and topical treatment. We found that topical treatment of this eyedrop markedly reduced corneal haze (Fig. 1D, E) and the expression of pro-inflammatory and pro-angiogenic genes at 3 days post NM injury when compared to vehicle- and NLE Ang (1-7) monomer-treated mice (Fig. 1F). The topical treatment-induced reduction of the expression of pro-inflammatory and pro-angiogenic genes was comparable to i.p. injection of NLE Ang (1-7) PLPs. These observations demonstrate the efficacy of topical treatment and i.p. injection of NLE Ang (1-7) PLPs in attenuating corneal inflammation at acute phase.

We also compared the anti-inflammatory effects of the NLE Ang (1-7) PLP to the topical corticosteroid Dexamethasone (4 mg/kg/day) which is one of the most commonly used anti-inflammatory medicines ^61,62^. Here, we found comparable decrease in cytokine expression between NLE Ang (1-7) PLP and Dexamethasone topical treatments (Fig. S9).

### Intra-peritoneal injection and topical treatment of NLE Ang (1-7) PLPs attenuate NM-induced corneal inflammation during delayed phase

To compare i.p. and topical routes over longer time periods, mice corneas were exposed to NM and treated daily for two weeks beginning the following day (Fig. 2). Mice (n=12) were either treated with saline only, NLE Ang (1-7) peptide (i.p. or topical, 0.5 mg/kg/day), or NLE Ang (1-7) PLP (i.p. or topical, 0.5 mg/kg/day) for 13 days (Fig. 2A). NLE Ang (1-7) PLP treatments significantly reversed the NM-induced expression of pro-inflammatory and pro-angiogenic genes in mouse corneas at 14 days post-injury (Fig. 2B).

**Figure 2.**
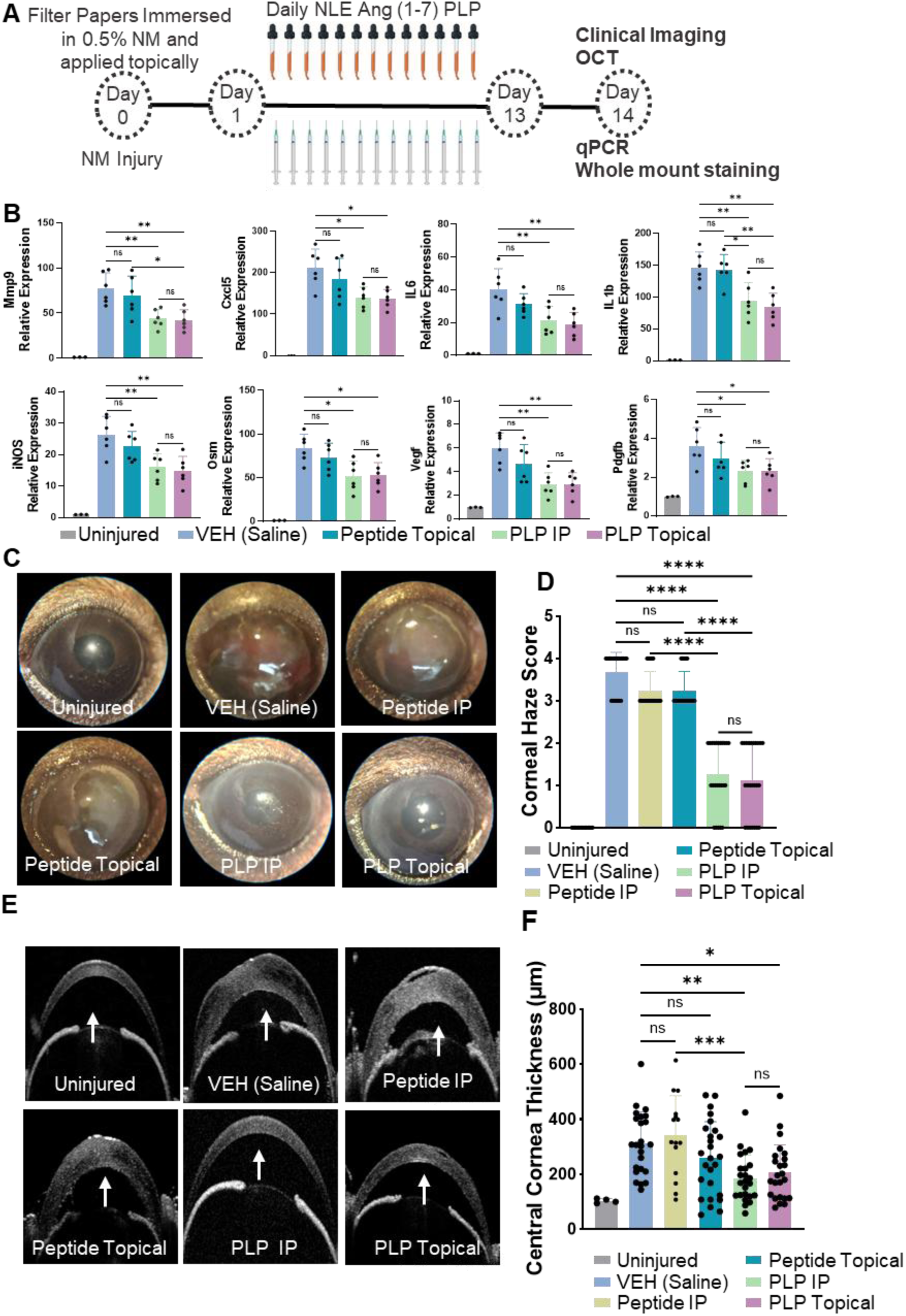
NLE Ang (1-7) PLP treatment attenuates corneal haze and corneal thickness in delayed phase of NM injury. (A) Schematic representation of 14 days treatment. Twenty-four hours post NM injury, mice were i.p injected or topically treated with NLE Ang (1-7) PLP or peptide (0.5 mg/kg/day) once daily for 13 days (delayed phase treatment). (B) At the end of treatment period, RT-qPCR was conducted to determine the expression of pro-inflammatory genes as well as genes related to angiogenesis in mouse cornea (n=6). (C) Slit lamp imaging was used to image the corneal haze. (D) Clinical examination of corneas revealed that NLE Ang(1-7) PLP treatments markedly decreased NM-induced corneal haze during delayed phase of NM injury (n=24). (E) Corneal thickness and structure were imaged using OCT. (F) OCT imaging of the corneas revealed that NLE Ang(1-7) PLP treatments markedly decreased NM-induced corneal thickening during delayed phase of NM injury (n=24) *: P ≤ 0.05, **:P ≤ 0.01, ***P ≤ 0.001, **** P ≤ 0.0001.

We examined the clinical indicators of corneal injury including corneal haze and edema. Mouse eyes were imaged on day 14 post NM exposure. Corneal haze scores were decreased in NLE Ang (1-7) PLP treated eyes (Fig. 2C, D). Optical coherence tomography (OCT) was also used as a non-invasive measurement of edema in mouse corneas and showed that NLE Ang (1-7) PLP treatments markedly diminished corneal thickening induced by NM exposure (Fig. 2E, F). More importantly, NLE Ang (1-7) PLP showed superior efficacy in reducing corneal haze and edema compared to NLE Ang (1-7) peptide using both i.p. and topical administration methods.

### NLE Ang (1-7) PLP prevents LSCD induced by NM exposure and resulting chronic inflammation during the delayed phase

It has been shown that exposure to mustard agents causes corneal neovascularization, a clinical sign of LSCD, and conjunctivalization (repopulation of the cornea with conjunctival epithelial and goblet cells), a hallmark of LSCD (Fig. 3) ^30,31,63–65^. To determine whether NLE Ang (1-7) PLP treatments can prevent neovascularization and conjunctivalization, at 14 days post-NM exposure, wholemounts of the entire cornea were stained with CD31 (an endothelial cell marker) to detect vessels (Figs. 3A-G) and MUC5AC (a goblet cell marker, Figs. 3H-N). NM exposure markedly induced corneal neovascularization with the detection of an increase in blood vessels in the central corneas (Fig. 3B). Goblet cells within the limbal/corneal epithelial area were absent in uninjured control mice (Fig. 3H) but readily detected in mice exposed to NM (Fig. 3I). NLE Ang (1-7) PLP treatment minimized the vascularized area in the cornea (Figs. 3 E-G) and markedly reduced goblet cells in the corneal epithelium (Figs. 3 L-N). Collectively, NLE Ang (1-7) PLP treatment showed significant efficacy in reverting neovascularization and conjunctivalization, characteristics of the delayed phase. In contrast, no therapeutic effect of NLE Ang (1-7) peptide treatment was detected (Figs. 3C, D, G, J, K, N).

**Figure 3.**
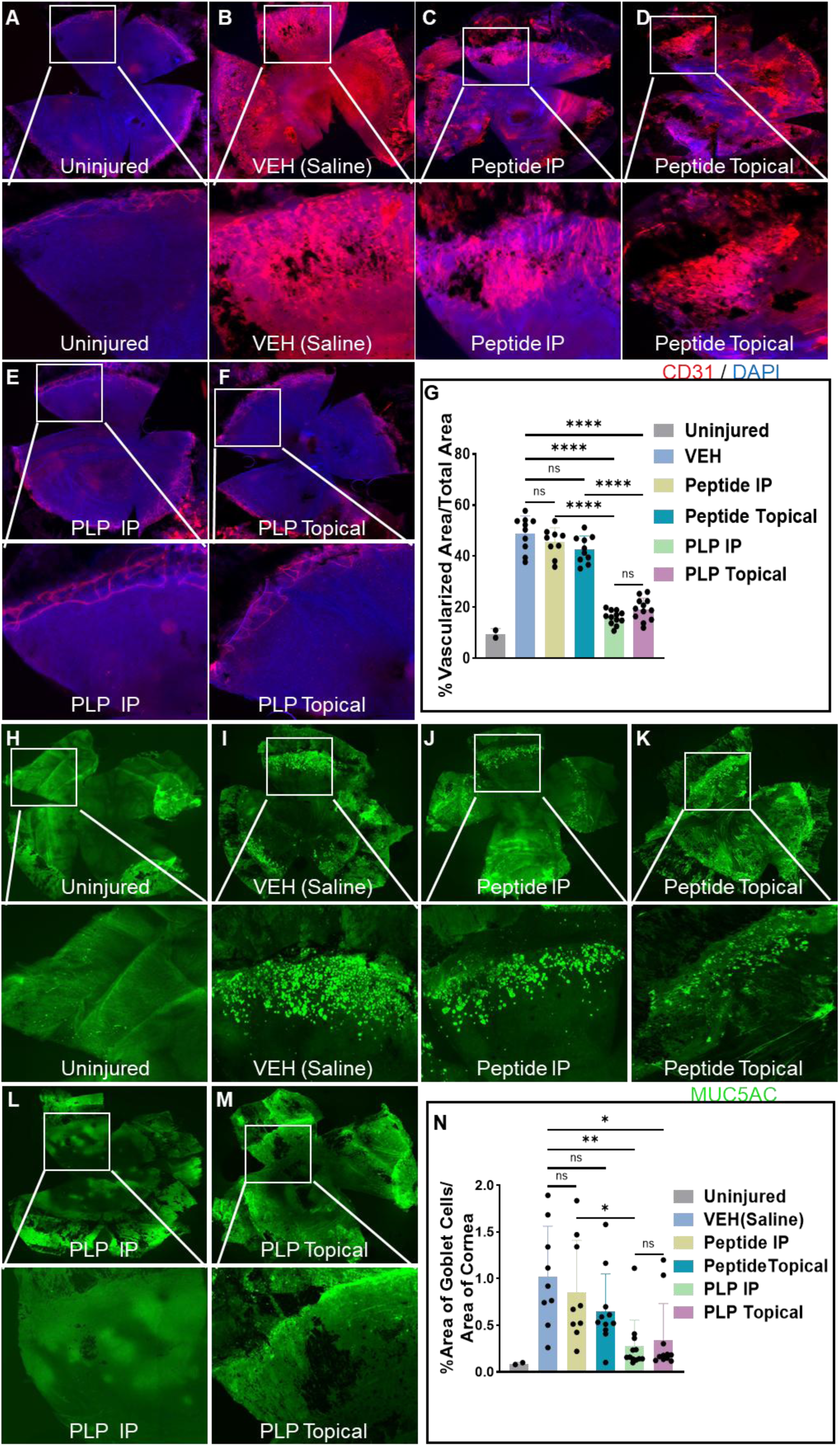
NLE Ang (1-7) PLP treatments attenuate vascularization and conjunctivalization in delayed phase of NM injury. (A-G) Whole mount immunostaining with CD31 demonstrated that PLP treatments (0.5 mg/kg/day) significantly reduced vascularization in the delayed phase of NM injury (n=10). Quantification was performed using ImageJ (G). The regions outlined by white boxes in the top panels are shown at higher magnification in the insets below. (H-N) Whole mount immunostaining with Mucin5AC (Muc5AC) revealed that PLP treatments (0.5 mg/kg/day) significantly reduced the area with goblet cells in cornea (n=10). Quantification was performed using ImageJ (N). The regions outlined by white boxes in the top panels are shown at higher magnification in the insets below. *: P ≤ 0.05, **:P ≤ 0.01, ***P ≤ 0.001, **** P ≤ 0.000

### Treatment of NM injury with NLE Ang (1-7) PLP shows persistent efficacy beyond the application period

We examined whether the therapeutic effects of the NLE Ang (1-7) PLP are sustained even after treatment stops (Fig. 4). One day post NM exposure, mice received once daily NLE Ang (1-7) PLP treatment (0.5 mg/kg/day) for 28 days, either by i.p. injection or topical eye drops (Fig. 4A). Eyes were scored for haze (Figs. 4B-H) and measured for edema (Figs. 4I-O) weekly. Four weeks after injury and daily treatments, the treatments stopped while the monitoring for corneal haze and edema continued for 28 more days followed by histology examination using H&E staining (Fig. 4P). NLE Ang (1-7) PLP-treated groups exhibited a significant reduction in corneal haze and edema compared to both vehicle- and monomeric peptide-treated controls over 56 days even after ceasing treatments. Remarkably, after treatment cessation, mice treated topically with PLP showed a continuous, significant improvement in measures of haze and edema (Figs. 4G, H, N, O). In contrast, corneal haze and edema were maintained in the mice treated with PLP i.p. after treatment stopped (Figs. 4F, H, M, O). Additionally, H&E staining showed a reduction of goblet cells in corneal epithelium and infiltrated cells in corneal stroma of both i.p. and topical NLE Ang (1-7) PLP treated mice (Fig. 4P). These observations demonstrate durable therapeutic benefits from early NLE Ang (1-7) PLP intervention.

**Figure 4.**
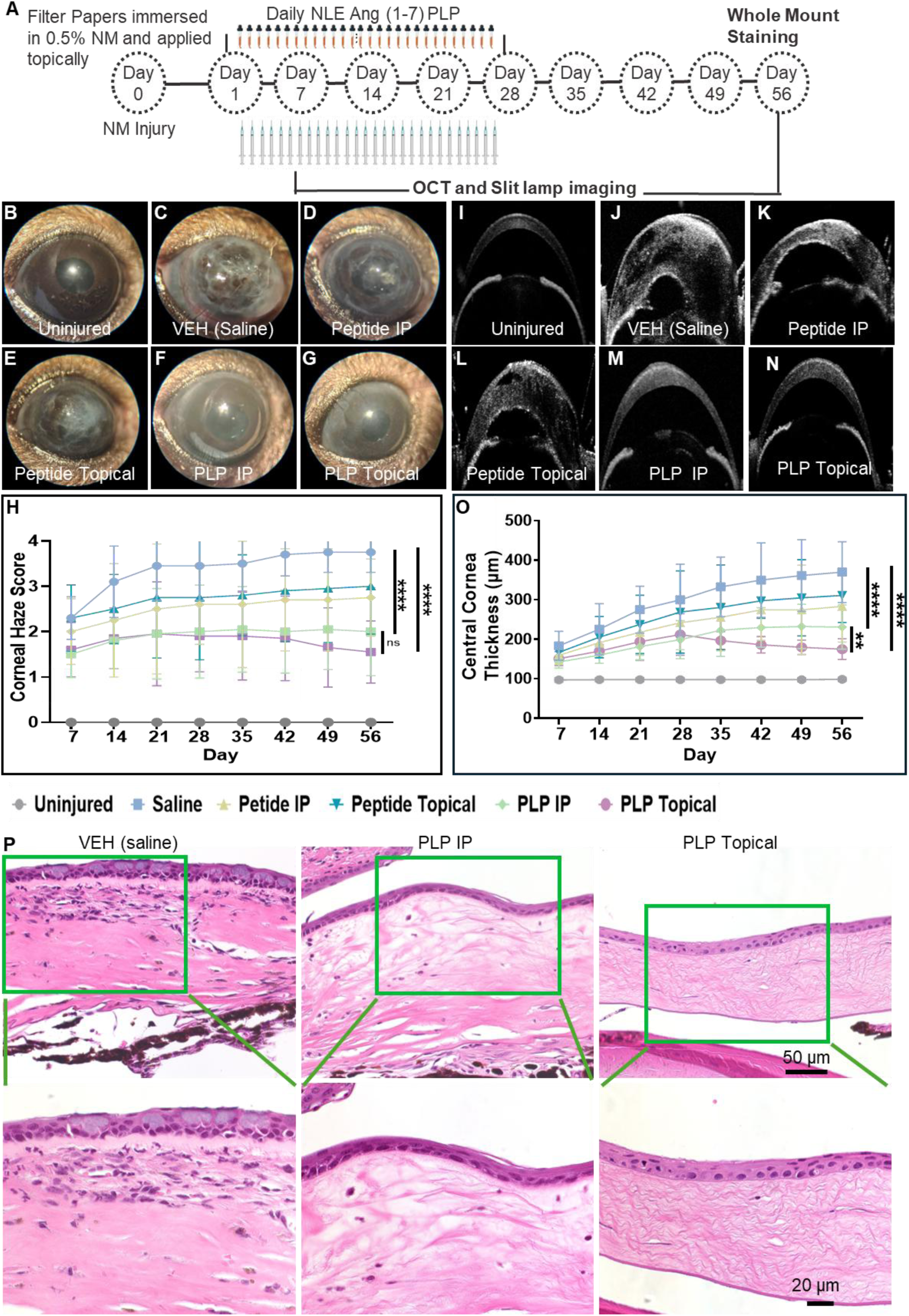
NLE Ang (1-7) PLP treatment reduces corneal haze, edema, inflammatory infiltration, and goblet cells in corneal epithelium over an extended post-injury period (56 days). (A) Schematic representation of 56 days experiment. Twenty-four hours post NM injury, mice were i.p. injected or topically treated with NLE Ang (1-7) PLP or NLE Ang (1-7) peptide (0.5 mg/kg/day) once daily for 28 days. On day 28, treatments were discontinued, and mice eyes were monitored weekly using slit-lamp and OCT machine until day 56. (B-G) Corneal haze and (I-O) OCT images for corneal thickness were imaged. (H) NLE Ang(1-7) PLP treatment markedly decreased NM-induced corneal haze even 28 days after treatment ceasing (n=20). (O) Corneal thickening was markedly decreased with NLE Ang(1-7) PLP treatment even 28 days after treatment ceasing (n=20). Quantification was conducted using ImageJ. *: P ≤ 0.05, **:P ≤ 0.01, ***P ≤ 0.001, **** P ≤ 0.0001. (P) H&E staining showed that PLP treatment reduced infiltration of immune cells, and goblet cell clusters in the cornea. The regions outlined by green boxes in the top panels are shown at higher magnification in the insets below.

### Topical application of NLE Ang (1-7) PLP precludes systemic exposure

Determination of systemic exposure and circulation of topically applied PLPs was assessed at several timepoints. Covalently labeled PLPs were synthesized for these studies using a Gadolinium (Gd) tag (see SI for synthetic details) ^56^. Here, the (Gd)-labeled PLPs were also formulated as eyedrops for application in the eyes of injured mice. Any detected Gd heavy metal in the organs or blood would be correlated to the PLP present in the sample as it is not naturally present in the body. Mice were harvested at timepoints of 48 hr., 24 hr., 8 hr., and 1 hr. post PLP treatment. Blood and liver samples were collected, digested, and run on ICP-MS. All samples showed that Gd levels were either below or detected within the noise of the instrument (LOD of 0.04 ppb, Table S3). This indicates that the PLP is not being detected in the systemic circulation following topical administration.

## Discussion and Conclusions

Topical therapy to prevent vision loss due to mustard injury requires targeting corneal epithelial and immune cells to expedite healing which we have achieved using proteinomimetic PLPs. Here, the time window between the initial corneal contact and subsequent immune activation is a feasible window for countermeasure intervention, allowing implementation of practical strategies in the event of mass emergency in pre-hospital triage and in hospitals ^66,67^. While the current clinical standards allow for the delivery of steroids to the eye, they cannot be used to treat the delayed phase of the injury due to their deleterious side effects ^34,35^.

The direct therapeutic use of Ang (1-7) peptides remains limited by significant pharmacologic and delivery challenges. As a small heptapeptide, Ang (1-7) is intrinsically unstable and undergoes rapid proteolytic degradation, resulting in extremely short half-life and poor bioavailability in ocular tissues as well as a demonstrated lack of efficacy in the model described here.

In response to these constraints and the clinical unmet need, several groups have turned to gene-based approaches such as adeno-associated virus (AAV)-mediated expression of Ang (1-7) to achieve sustained intraocular production, circumventing the peptide’s instability and enabling durable activation of the protective RAS axis ^38^. However, the clinical translation of AAV vectors remains constrained by several major challenges including immune responses against AAV capsid, dose dependent toxicity, limited packaging capacity, concerns about long term safety and carcinogenicity in high dose ^68,69^. In this work, we show the efficacy of the NLE Ang (1-7) peptide delivered both topically and systemically. Most importantly, the NLE Ang (1-7) proteomimetic polymer demonstrated extended enzyme stability, superior efficacy and continued anti-inflammatory effects even after treatment cessation. Additionally, the proteomimetic polymer exhibited superior efficacy in the less invasive topical application over systemic injections. This is of particular interest because of the ease of use and increased patient compliance of eyedrops compared to injections. This instability compounded by the structural and enzymatic barriers of the eye including the tear film, corneal epithelium, and local proteases also severely restrict peptide penetration and retention of other peptide and protein therapeutics, making topical or intravitreal delivery inefficient. These limitations are consistent with the broader challenges faced by peptide and protein therapeutics across many diseases ^70^. The PLP platform technology can address these barriers by providing in vivo stability, potent intracellular bioactivity, and multivalency, enabling efficient utilization of diverse peptides in multiple disease contexts ^54–58^. For example, we have demonstrated that PLP conjugated HV3 peptides show a 2000-fold increase in circulation half-life compared to peptides in vivo and selectively target mutant huntingtin in Huntington’s disease (HD) models ^56^. In a rat myocardial infarction model, Keap1-binding peptide delivered via PLP improved cardiac function through immunomodulatory, anti-apoptotic, and angiogenic mechanisms ^55^.

PLP shows similar effects to the current steroid (Dexamethasone) that is administered in the clinic. However, the steroid can only be administered for short periods of time due to the severe side effects of long term use ^71–73^. Long term use is needed because, following resolution of the acute corneal mustard injury, a delayed phase can occur in patients after months or even years of a clinically silent period ^74^. Importantly, the superior efficacy of the NLE Ang (1-7) PLP in alleviating mustard-induced corneal injury can persist for over two months in our mouse model, which approximates a 6-year duration in humans ^75,76^. This highlights the potential of the PLP in providing a long term therapeutic benefit.

In summary, we have described an angiotensin-inspired PLP as being effective at decreasing acute and delayed inflammation in the corneal following nitrogen mustard injuries. Based on these findings, these proteomimetic polymers have potential in meeting an unmet clinical need in the treatment of chemical injury in the eye for both acute and prolonged inflammation. Ongoing work involves investigating other corneal diseases and injuries that would benefit from topical delivery of PLPs as eye drops.

## Acknowledgments

Imaging was performed at the Northwestern University Center for Advanced Microscopy generously supported by NCI CCSG P30 CA060553 awarded to the Robert H. Lurie Comprehensive Cancer Center. This research was supported in part by resources provided by the Northwestern University Skin Biology and Diseases Resource-based Center (NIAMS P30AR075049). This research was also supported by National Institutes of Health grant EY019463 (RML), National Institutes of Health grants EY032922, EY028560, and EY037653 (HP). This research and N.C.G. are supported by National Science Foundation for the PLP platform technology grant DMR-2403954 and National Institutes of Health grant 706200 AMND 2//5R01HL139001-07.

## Author Contributions

P.F., A.K.N., R.L., H.P. and N.C.G. conceived and designed research. A.K.N, M.T and J.R.M. conducted the synthesis and characterization of materials. J.R.M. and A.K.N. performed enzyme degradation assays. A.F. performed cellular uptake. P.F. and H.P. performed *in vivo* NM, RT-qPCR, and vascularization assessment. H.P., R.L. and N.C.G. supervised the project. All authors discussed the results and contributed to writing the manuscript.

## Data Availability Statement

All relevant data supporting the findings of this study are available within the paper and Supplementary Information.

## Ethics approval statement

All procedures received approval from the Institutional Animal Care and Use Committee (IACUC) at Northwestern University and are in adherence to the ARVO Statement for the Use of Animals in Ophthalmic and Vision Research.

## References

1 Basu, S. et al. Human limbal biopsy-derived stromal stem cells prevent corneal scarring. Sci Transl Med 6, 266ra172, doi:10.1126/scitranslmed.3009644 (2014).

2 Cotsarelis, G., Cheng, S. Z., Dong, G., Sun, T. T. & Lavker, R. M. Existence of slow-cycling limbal epithelial basal cells that can be preferentially stimulated to proliferate: implications on epithelial stem cells. Cell 57, 201–209 (1989).

3 Funderburgh, J. L., Funderburgh, M. L. & Du, Y. Stem Cells in the Limbal Stroma. Ocul Surf 14, 113–120, doi:10.1016/j.jtos.2015.12.006 (2016).

4 Lehrer, M. S., Sun, T. T. & Lavker, R. M. Strategies of epithelial repair: modulation of stem cell and transit amplifying cell proliferation. Journal of cell science 111 **(Pt** **19**), 2867–2875 (1998).

5 Pajoohesh-Ganji, A. & Stepp, M. A. In search of markers for the stem cells of the corneal epithelium. Biol Cell 97, 265–276, doi:10.1042/BC20040114 (2005).

6 Parfitt, G. J. et al. Immunofluorescence Tomography of Mouse Ocular Surface Epithelial Stem Cells and Their Niche Microenvironment. Invest Ophthalmol Vis Sci 56, 7338–7344, doi:10.1167/iovs.15-18038 (2015).

7 Peng, H. et al. MicroRNA-31 targets FIH-1 to positively regulate corneal epithelial glycogen metabolism. FASEB J 26, 3140–3147, doi:10.1096/fj.11-198515 (2012).

8 Stepp, M. A. & Zieske, J. D. The corneal epithelial stem cell niche. Ocul Surf 3, 15–26 (2005).

9 Schermer, A., Galvin, S. & Sun, T. T. Differentiation-related expression of a major 64K corneal keratin in vivo and in culture suggests limbal location of corneal epithelial stem cells. J Cell Biol 103, 49–62, doi:10.1083/jcb.103.1.49 (1986).

10 Imanishi, J. et al. Growth factors: importance in wound healing and maintenance of transparency of the cornea. Prog Retin Eye Res 19, 113–129, doi:10.1016/s1350-9462(99)00007-5 (2000).

11 Sotozono, C. et al. Cytokine expression in the alkali-burned cornea. Curr Eye Res 16, 670–676, doi:10.1076/ceyr.16.7.670.5057 (1997).

12 Burns, A. R., Li, Z. & Smith, C. W. Neutrophil migration in the wounded cornea: the role of the keratocyte. Ocul Surf 3, S173–176, doi:10.1016/s1542-0124(12)70249-5 (2005).

13 Pfister, R. R. & Haddox, J. L. A neutrophil chemoattractant is released from cellular and extracellular components of the alkali-degraded cornea and blood. Invest Ophthalmol Vis Sci 37, 230–237 (1996).

14 Saika, S. Yin and yang in cytokine regulation of corneal wound healing: roles of TNF-alpha. Cornea 26, S70–74, doi:10.1097/ICO.0b013e31812f6d14 (2007).

15 Jin, Y. et al. The chemokine receptor CCR7 mediates corneal antigen-presenting cell trafficking. Mol Vis 13, 626–634 (2007).

16 Spandau, U. H., Toksoy, A., Verhaart, S., Gillitzer, R. & Kruse, F. E. High expression of chemokines Gro-alpha (CXCL-1), IL-8 (CXCL-8), and MCP-1 (CCL-2) in inflamed human corneas in vivo. Arch Ophthalmol 121, 825–831, doi:10.1001/archopht.121.6.825 (2003).

17 Wallace, G. R., John Curnow, S., Wloka, K., Salmon, M. & Murray, P. I. The role of chemokines and their receptors in ocular disease. Prog Retin Eye Res 23, 435–448, doi:10.1016/j.preteyeres.2004.04.004 (2004).

18 Kayaalp Nalbant, E., et al. A novel therapy to ameliorate nitrogen mustard-induced limbal stem cell deficiency using lipoprotein-like nanoparticles. NPJ Regen Med 10, 14, doi:10.1038/s41536-025-00402-5 (2025).

19 Abdelfattah, N. S., Amgad, M., Zayed, A. A., Hussein, H. & Abd El-Baky, N. Molecular underpinnings of corneal angiogenesis: advances over the past decade. Int J Ophthalmol 9, 768–779, doi:10.18240/ijo.2016.05.24 (2016).

20. Azar, D. T. Corneal angiogenic privilege: angiogenic and antiangiogenic factors in corneal avascularity, vasculogenesis, and wound healing (an American Ophthalmological Society thesis). Trans Am Ophthalmol Soc 104, 264–302 (2006).

21 Clements, J. L. & Dana, R. Inflammatory corneal neovascularization: etiopathogenesis. Semin Ophthalmol 26, 235–245, doi:10.3109/08820538.2011.588652 (2011).

22 Koo, H. Y. & Kume, T. FoxC1-dependent regulation of vascular endothelial growth factor signaling in corneal avascularity. Trends Cardiovasc Med 23, 1–4, doi:10.1016/j.tcm.2012.08.002 (2013).

23 Notara, M., Lentzsch, A., Coroneo, M. & Cursiefen, C. The Role of Limbal Epithelial Stem Cells in Regulating Corneal (Lymph)angiogenic Privilege and the Micromilieu of the Limbal Niche following UV Exposure. Stem Cells Int 2018, 8620172, doi:10.1155/2018/8620172 (2018).

24 Rafati-Rahimzadeh, M. et al. Ophthalmological aspects of mustard gas poisoning (focus on management). Caspian J Intern Med 13, 458–468, doi:10.22088/cjim.13.3.458 (2022).

25 Aasted, A., Darre, E. & Wulf, H. C. Mustard gas: clinical, toxicological, and mutagenic aspects based on modern experience. Ann Plast Surg 19, 330–333 (1987).

26 Baradaran-Rafii, A., Eslani, M. & Tseng, S. C. Sulfur mustard-induced ocular surface disorders. Ocul Surf 9, 163–178, doi:10.1016/s1542-0124(11)70026-x (2011).

27 Panahi, Y., Rajaee, S. M. & Sahebkar, A. Ocular Effects of Sulfur Mustard and Therapeutic Approaches. J Cell Biochem 118, 3549–3560, doi:10.1002/jcb.25892 (2017).

28 Saladi, R. N., Smith, E. & Persaud, A. N. Mustard: a potential agent of chemical warfare and terrorism. Clin Exp Dermatol 31, 1–5, doi:10.1111/j.1365-2230.2005.01945.x (2006).

29 Kadar, T. et al. Characterization of acute and delayed ocular lesions induced by sulfur mustard in rabbits. Curr Eye Res 22, 42–53, doi:10.1076/ceyr.22.1.42.6975 (2001).

30 McNutt, P. M. et al. Contributions of tissue-specific pathologies to corneal injuries following exposure to SM vapor. Ann N Y Acad Sci 1374, 132–143, doi:10.1111/nyas.13105 (2016).

31 Safarinejad, M. R., Moosavi, S. A. & Montazeri, B. Ocular injuries caused by mustard gas: diagnosis, treatment, and medical defense. Mil Med 166, 67–70 (2001).

32 Fuchs, A., Giuliano, E. A., Sinha, N. R. & Mohan, R. R. Ocular toxicity of mustard gas: A concise review. Toxicol Lett 343, 21–27, doi:10.1016/j.toxlet.2021.02.007 (2021).

33 Kadar, T. et al. Prolonged impairment of corneal innervation after exposure to sulfur mustard and its relation to the development of delayed limbal stem cell deficiency. Cornea 32, e44–50, doi:10.1097/ICO.0b013e318262e885 (2013).

34 Petroutsos, G., Guimaraes, R., Giraud, J. P. & Pouliquen, Y. Corticosteroids and corneal epithelial wound healing. Br J Ophthalmol 66, 705–708, doi:10.1136/bjo.66.11.705 (1982).

35 Balali-Mood, M. & Hefazi, M. The pharmacology, toxicology, and medical treatment of sulphur mustard poisoning. Fundam Clin Pharmacol 19, 297–315, doi:10.1111/j.1472-8206.2005.00325.x (2005).

36 Abdallah, W. F. et al. NorLeu3A(1-7) Accelerates Clear Corneal Full Thickness Wound Healing. Invest Ophthalmol Vis Sci 57, 2187–2194, doi:10.1167/iovs.15-18515 (2016).

37 Wang, J. et al. The ACE2-deficient mouse: A model for a cytokine storm-driven inflammation. Faseb j 34, 10505–10515, doi:10.1096/fj.202001020R (2020).

38 Verma, A. et al. ACE2 and Ang-(1-7) confer protection against development of diabetic retinopathy. Mol Ther 20, 28–36, doi:10.1038/mt.2011.155 (2012).

39 Giese, M. J. & Speth, R. C. The ocular renin-angiotensin system: a therapeutic target for the treatment of ocular disease. Pharmacol Ther 142, 11–32, doi:10.1016/j.pharmthera.2013.11.002 (2014).

40 Jadhav, S. S. et al. Effects of combined radiation and burn injury on the renin-angiotensin system. Wound Repair Regen 21, 131–140, doi:10.1111/j.1524-475X.2012.00867.x (2013).

41 Durik, M., Seva Pessoa, B. & Roks, A. J. The renin-angiotensin system, bone marrow and progenitor cells. Clin Sci (Lond) 123, 205–223, doi:10.1042/CS20110660 (2012).

42 Rodgers, K. E., Bolton, L. L., Verco, S. & diZerega, G. S. NorLeu(3)-Angiotensin (1-7) [DSC127] as a Therapy for the Healing of Diabetic Foot Ulcers. Adv Wound Care (New Rochelle) 4, 339–345, doi:10.1089/wound.2014.0609 (2015).

43 Passos-Silva, D. G., Verano-Braga, T. & Santos, R. A. Angiotensin-(1-7): beyond the cardio-renal actions. Clin Sci (Lond) 124, 443–456, doi:10.1042/CS20120461 (2013).

44 Rodgers, K. E. & Dizerega, G. S. Contribution of the Local RAS to Hematopoietic Function: A Novel Therapeutic Target. Front Endocrinol (Lausanne) 4, 157, doi:10.3389/fendo.2013.00157 (2013).

45 Rodgers, K. E. et al. Effect of NorLeu3-A(1-7) on scar formation over time after full-thickness incision injury in the rat. Wound Repair Regen 13, 309–317, doi:10.1111/j.1067-1927.2005.130314.x (2005).

46 Rodgers, K. E. et al. Acceleration of healing, reduction of fibrotic scar, and normalization of tissue architecture by an angiotensin analogue, NorLeu3-A(1-7). Plast Reconstr Surg 111, 1195–1206, doi:10.1097/01.PRS.0000047403.23105.66 (2003).

47 Rodgers, K. et al. Development of angiotensin (1-7) as an agent to accelerate dermal repair. Wound Repair Regen 9, 238–247, doi:10.1046/j.1524-475x.2001.00238.x (2001).

48 Rodgers, K., Verco, S., Bolton, L. & Dizerega, G. Accelerated healing of diabetic wounds by NorLeu(3)-angiotensin (1-7). Expert Opin Investig Drugs 20, 1575–1581, doi:10.1517/13543784.2011.619976 (2011).

49 Blum, A. P. et al. Peptides displayed as high density brush polymers resist proteolysis and retain bioactivity. J Am Chem Soc 136, 15422–15437, doi:10.1021/ja5088216 (2014).

50 Bielawski, C. W. & Grubbs, R. H. Living ring-opening metathesis polymerization. Prog Polym Sci 32, 1–29, doi:10.1016/j.progpolymsci.2006.08.006 (2007).

51 Blum, A. P. et al. Stimuli Induced Uptake of Protein-Like Peptide Brush Polymers. Chemistry 28, e202103438, doi:10.1002/chem.202103438 (2022).

52 Blum, A. P., Kammeyer, J. K. & Gianneschi, N. C. Activating Peptides for Cellular Uptake via Polymerization into High Density Brushes. Chem Sci 7, 989–994, doi:10.1039/C5SC03417E (2016).

53 Sun, H. et al. Origin of Proteolytic Stability of Peptide-Brush Polymers as Globular Proteomimetics. ACS Cent Sci 7, 2063–2072, doi:10.1021/acscentsci.1c01149 (2021).

54 Carrow, K. P. et al. Inhibiting the Keap1/Nrf2 Protein-Protein Interaction with Protein-Like Polymers. Adv Mater 36, e2311467, doi:10.1002/adma.202311467 (2024).

55 Mesfin, J. M. et al. Protein-Like Polymers Targeting Keap1/Nrf2 as Therapeutics for Myocardial Infarction. Advanced Materials 37, e2417885, doi:ARTN 2417885 10.1002/adma.202417885 (2025).

56 Choi, W. et al. Proteomimetic polymer blocks mitochondrial damage, rescues Huntington’s neurons, and slows onset of neuropathology in vivo. Sci Adv 10, eado8307, doi:10.1126/sciadv.ado8307 (2024).

57 Berger, O. et al. Mussel Adhesive-Inspired Proteomimetic Polymer. J Am Chem Soc 144, 4383–4392, doi:10.1021/jacs.1c10936 (2022).

58 Choi, W. et al. Thrombospondin-1 proteomimetic polymers exhibit anti-angiogenic activity in a neovascular age-related macular degeneration mouse model. Sci Adv 9, eadi8534, doi:10.1126/sciadv.adi8534 (2023).

59 Merrifield, R. B. Solid Phase Peptide Synthesis. I. The Synthesis of a Tetrapeptide. Journal of the American Chemical Society 85, 2149–2154 (1963).

60 Coin, I., Beyermann, M. & Bienert, M. Solid-phase peptide synthesis: from standard procedures to the synthesis of difficult sequences. Nat Protoc 2, 3247–3256, doi:10.1038/nprot.2007.454 (2007).

61 Abadia, B. et al. Clinical Applications of Dexamethasone for Aged Eyes. Drugs Aging 33, 639–646, doi:10.1007/s40266-016-0392-z (2016).

62 Gaballa, S. A. et al. Corticosteroids in ophthalmology: drug delivery innovations, pharmacology, clinical applications, and future perspectives. Drug Deliv Transl Res 11, 866–893, doi:10.1007/s13346-020-00843-z (2021).

63 Le, Q., Xu, J. & Deng, S. X. The diagnosis of limbal stem cell deficiency. Ocul Surf 16, 58–69, doi:10.1016/j.jtos.2017.11.002 (2018).

64 Javadi, M. A. et al. Chronic and delayed-onset mustard gas keratitis: report of 48 patients and review of literature. Ophthalmology 112, 617–625, doi:10.1016/j.ophtha.2004.09.027 (2005).

65 Lavker, R. M., Tseng, S. C. G. & Sun, T. T. Corneal epithelial stem cells at the limbus: looking at some old problems from a new angle. Experimental Eye Research 78, 433–446, doi:10.1016/j.exer.2003.09.008 (2004).

66 Goswami, D. G. et al. Pathophysiology and inflammatory biomarkers of sulfur mustard-induced corneal injury in rabbits. PLoS One 16, e0258503, doi:10.1371/journal.pone.0258503 (2021).

67 McNutt, P. Progress towards a standardized model of ocular sulfur mustard injury for therapeutic testing. Exp Eye Res 228, 109395, doi:10.1016/j.exer.2023.109395 (2023).

68 Colella, P., Ronzitti, G. & Mingozzi, F. Emerging Issues in AAV-Mediated In Vivo Gene Therapy. Mol Ther Methods Clin Dev 8, 87–104, doi:10.1016/j.omtm.2017.11.007 (2018).

69 Moldavskii, D. et al. AAV-Based Gene Therapy: Opportunities, Risks, and Scale-Up Strategies. Int J Mol Sci 26, doi:10.3390/ijms26178282 (2025).

70 Wang, L. et al. Therapeutic peptides: current applications and future directions. Signal Transduct Target Ther 7, 48, doi:10.1038/s41392-022-00904-4 (2022).

71 Fraunfelder, F. W. Corneal toxicity from topical ocular and systemic medications. Cornea 25, 1133–1138, doi:10.1097/01.ico.0000240084.27663.fd (2006).

72 Holland, E. J., Fingeret, M. & Mah, F. S. Use of Topical Steroids in Conjunctivitis: A Review of the Evidence. Cornea 38, 1062–1067, doi:10.1097/ico.0000000000001982 (2019).

73 Musleh, M. G., Bokre, D. & Dahlmann-Noor, A. H. Risk of intraocular pressure elevation after topical steroids in children and adults: A systematic review. Eur J Ophthalmol 30, 856–866, doi:10.1177/1120672119885050 (2020).

74 Kadar, T. et al. Ocular injuries following sulfur mustard exposure--pathological mechanism and potential therapy. Toxicology 263, 59–69, doi:10.1016/j.tox.2008.10.026 (2009).

75 Cottam, N. C. et al. From Circuits to Lifespan: Translating Mouse and Human Timelines with Neuroimaging-Based Tractography. J Neurosci 45, e1429242025, doi:10.1523/JNEUROSCI.1429-24.2025 (2025).

76 The Jackson Laboratory. When are mice considered old?, & <https://www.jax.org/news-and-insights/jax-blog/2017/november/when-are-mice-considered-old> (2017).

