## Supplemental Methods and Materials, Figures and Tables for "Angiotensin (1-7)-based Proteomimetic Polymers as Topicals for the Remediation and Repair of the Cornea following Injury"

**SUPPLEMENTAL INFORMATION:**

**TABLE OF CONTENTS**

**Materials and Methods ……………………………………………………………..………. 3**

I. **Materials …………………………………………………………………………... 3**
II. **Instrumentation ……………………………….……………………….……...…. 3**
III. **Experimental Methods ……………………………………….……….…...……. 4**

**List of Figures ………………………………………………………………………….……. 10**

**Supplemental Figures ……………………..……………………..……………….......…… 11**

**Supplemental Tables ……………………………………………....………………………. 17**

**References ………………………………………………………….…………...……..……. 20**

**Materials and Methods**

I. ***Materials***

All materials and reagents, unless otherwise noted, were purchased from commercial sources, and used as received. Abbreviations of L-amino acids are denoted by uppercase. *N*-(hexanoic acid)-*cis*-5-norbornene-*exo*-dicarboximide, *N*-(pentanol)-*cis*-5-norbornene-*exo*-dicarboximide and the olefin metathesis initiator (IMesH_2_)(C_5_H_5_N)_2_(Cl)_2_Ru=CHPh were synthesized according to our previous report (1–3). Protein gels (Any kD Mini-PROTEAN, 10 well) were purchased from BioRad (catalog No. 4569033). Dulbecco’s Phosphate Buffered Saline (DPBS, catalog No. 45000-434) and Fetal Bovine Serum (FBS, heat inactivated, catalog No. 35011CV) were purchased from Corning. Antibiotics (Penicillin-Streptomycin, catalog No. 15140122) were purchased from Life Technologies. Human corneal epithelial cell line were generously donated from the Jester Lab (UCI Gavin Herbert Eye Institute. Cell media (Keratinocyte SFM 1x) was purchased from ThermoFischer Scientific (catalog No. 17005042). The mice primer pairs for RT‐qPCR were purchased from IDT (Coralville, IA, USA). Trypsin enzyme (Platinum grade, catalog No. VA9000) for stability assays were purchased from Promega Corporation.

II. ***Instrumentation***
i. **Solid Phase Peptide Synthesis (SPPS):** Peptides were synthesized using standard Fmoc-chemistry SPPS procedures on either an AAPPTec Focus XC automated synthesizer or on a Liberty Blue (CEM) automated microwave peptide synthesizer.
ii. **^1^H Nuclear Magnetic Resonance (^1^H NMR):** ^1^H NMR spectra were recorded on one of the following: Varian Inova spectrometer (500 MHz), Bruker Avance III HD Nanobay (400 MHz), Bruker Avance III HD system (500 MHz), and Bruker Avance III (600 MHz). Chemical shifts (^1^H) are reported in (ppm) relative to the residual solvent peak.
iii. **Analytical High-Performance Liquid Chromatography (Analytical HPLC):** Analytical analysis of peptides and peptide monomers was performed on a Jupiter 4 Proteo 90Å Phenomenex column (150 x 4.60 mm) using a Hitachi-Elite LaChrom L-2130 pump equipped with UV-Vis detector (Hitachi-Elite LaChrom L2420). The solvent system consists of (A) 0.1% TFA in 99.9% water and (B) 0.1% TFA in 99.9% acetonitrile.
iv. **Automated Flash Chromatography:** Peptide monomers were purified using a Biotage Selekt flash chromatography system. The solvent system consists of (A) 0.1% TFA in water and (B) 0.1% TFA in acetonitrile.
v. **Preparative HPLC:** An Armen Glider CPC preparatory HPLC and a BUCHI Pure C-830 Preparative HPLC was used to purify all peptides and peptide monomers. The solvent system consists of (A) 0.1% TFA in water and (B) 0.1% TFA in acetonitrile.
vi. **Electrospray Ionization Mass Spectrometry:** ESI-MS spectra of peptides and peptide monomers were collected using a Bruker Amazon-SL spectrometer configured with an ESI source in both negative and positive ionization mode.
vii. **Mass Assisted Laser Desorption Ionization-Time of Flight:** MALDI-MS spectra were collected using a Bruker rapileX Tissue typer configured with a laser source (1000-5000 shots, 1 kHz, laser beam attenuation 60-40%) in negative and positive ion mode.
viii. **Dynamic Light Scattering:** Dynamic light scattering was performed using a Wyatt DynaPro NanoStar at 657 nm.
ix. **Fluorescence Measurement:** CellTiter-Blue® fluorescence measurements were recorded using a Perkin Elmer EnSpire multimode Plate Reader at 590 nm.
x. **Animal Studies:** All animal studies were conducted in accordance with NIH guidelines for the care and use of laboratory animals and protocols were approved by the Institutional Animal Care and Use Committee of Northwestern University. Six to eight-week-old C57BL/6J female mice were purchased from Jackson Laboratories.
xi. **ICP-MS:** Inductively couple plasma mass spectrometry (ICP-MS) was performed at the Northwestern University Quantitative Bioelemental Imaging Core on the Thermo iCAP Q ICP-MS, controlled using QTEGRA software.
III. ***Experimental Methods***
i. **Preparation of peptide monomers via solid phase peptide synthesis (SPPS):** All peptide monomers were synthesized on Protide resin using standard Fmoc SPPS procedures on an AAPPTec Focus XC automated synthesizer or Liberty Blue Automated Microwave Synthesizer. Peptide monomers connected via amide bond on the C-terminus (**M1**, **M3**, and **M1Scr**) were prepared via amide coupling to N-(hexanoic acid)-cis-5-norbornene-exo-dicarboximide (3.0 equiv) in the presence of HBTU (2.9 equiv) and DIPEA (6.0 equiv). The peptide monomers were cleaved from resin by treatment with TFA/H_2_O/TIPS (95:2.5:2.5 v/v) for 4 hr. The crude products were filtered to remove the resin and precipitated in cold diethyl ether. The pellet was dried under reduced pressure and purified via automated flash chromatography and preparative HPLC.
ii. **General procedure for NMR-scale polymerization reaction kinetics:** Polymerizations were performed under dry dinitrogen atmosphere. Norbornene conjugated peptide monomers (25 mg/mL, 15.0 equiv.) were dissolved in DMF-*d_7_* and an initial NMR experiment was conducted. Grubbs’ catalyst (IMesH_2_)(C_5_H_5_N)_2_(Cl)_2_ Ru=CHPh (1.0 equiv., 20 mg/mL in DMF) was added and the reaction was tracked via NMR until full conversion of the monomer. The reaction was quenched with ethyl vinyl ether (EVE, excess) and stirred for 30 mins. The polymer solution was precipitated into diethyl ether and centrifuged. The product was directly injected with preparative HPLC. The product was collected and dried by lyophilization as a white solid.
iii. **General procedure for ring opening metathesis polymerization (ROMP) of homopolymer PLPs:** Polymerizations were performed under dry dinitrogen atmosphere. Norbornene conjugated peptide monomers (25 mg/mL, 15.0 equiv.) were dissolved in degassed DMF. Grubbs’ catalyst (IMesH_2_)(C_5_H_5_N)_2_(Cl)_2_ Ru=CHPh (1.0 equiv., 20 mg/mL in DMF) was added and the reaction was stirred for 2-5 hr. After full conversion of the monomer, the reaction was quenched with ethyl vinyl ether (EVE, excess) and stirred for 30 mins. The product was directly injected with preparative HPLC. The product was collected and dried by lyophilization as a white solid.

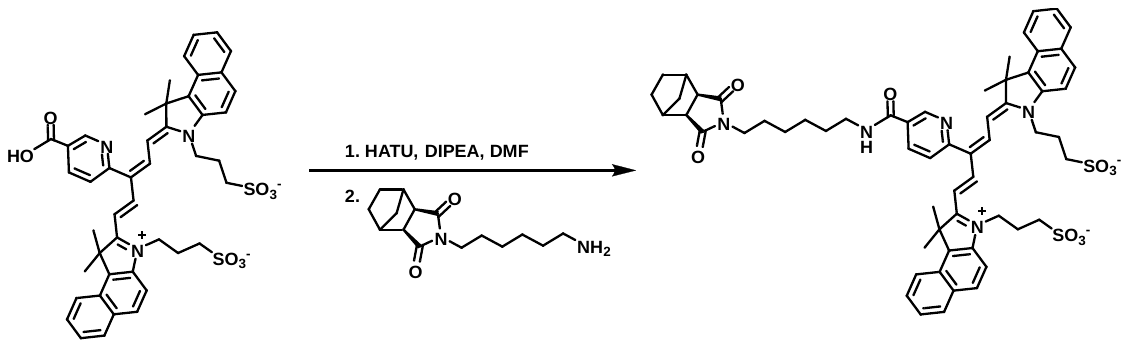

iv. **Synthesis of Cy5 monomer (Nor-Cy5)** (4)**:** To cyanine (75 mg, 0.091 mmol) in dimethylformamide (1.5 mL) was added DIPEA (192 µl, 1.1 mmol) and HATU (34.8 mg, 0.091 mmol). The reaction was subsequently stirred for 10 min, norbornene amine (66 mg, 0.251 mmol) was added, and stirred for 3 hours. The reaction was precipitated into cold diethyl ether, centrifuged, and the precipitate was dried. The crude product was purified by flash chromatography (35-55% ethyl acetate: hexanes) resulting in a blue solid. ESI-MS expected [M+H]^+^: 1064.43, found 1064.52. NMR spectrum was run in CD_3_OD.

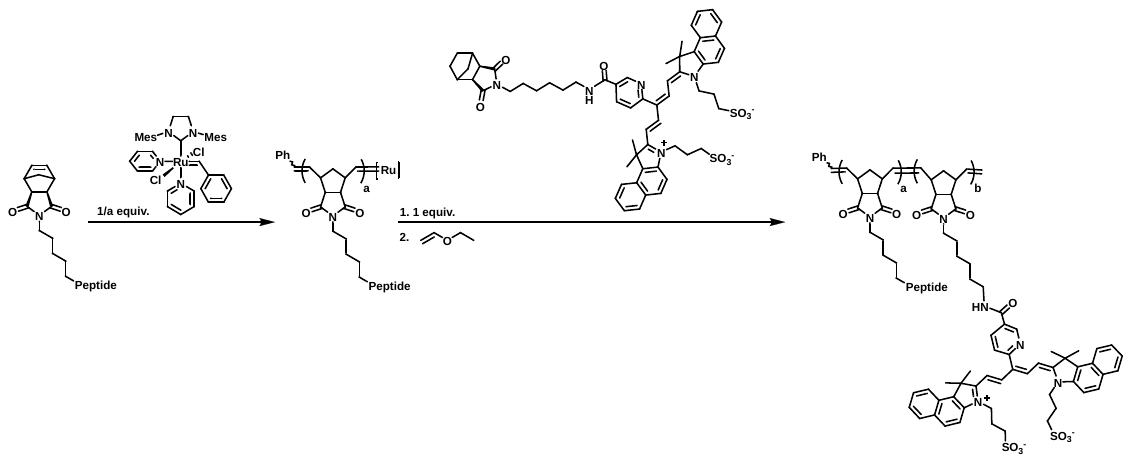

v. **Synthesis of** **NLE Ang (1-7) PLP-Cy5:** Cyanine addition was performed as previously reported (5). Polymerizations were performed under dry dinitrogen atmosphere. **M1** (25 mg/mL, n equiv.) was dissolved in degassed DMF. Grubbs’ catalyst (IMesH_2_)(C_5_H_5_N)_2_(Cl)_2_Ru=CHPh (1/n equiv., 20 mg/mL in DMF) was added and the reaction was stirred for 3 hours. **Nor-Cy5** (1 equiv.) was added and the reaction stirred for 2 hours. The polymerization was terminated with EVE (excess) and stirred for 30 mins. The polymer solution was precipitated into diethyl ether and centrifuged. The pellet was dried under reduced pressure and dialyzed in MilliQ water. The resulting solution was lyophilized resulting in a blue solid.

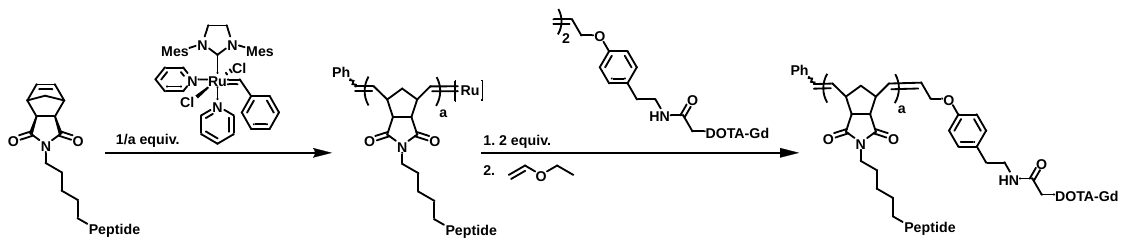

vi. **Synthesis of NLE Ang (1-7) PLP-Gd:** Terminations were performed as previously reported(6). Polymerizations were performed under dry dinitrogen atmosphere. **M1** (25 mg/mL, n equiv.) was dissolved in degassed DMF. Grubbs’ catalyst (IMesH_2_)(C_5_H_5_N)_2_(Cl)_2_Ru=CHPh (1/n equiv., 20 mg/mL in DMF) was added and the reaction was stirred for 3 hours. **Gd-TA** (2 equiv.) was added and the reaction stirred for 6 hours. The polymerization was terminated with EVE (excess) and stirred for 30 mins. The polymer solution was precipitated into diethyl ether and centrifuged. The pellet was dried under reduced pressure and dialyzed in MilliQ water. The resulting solution was lyophilized resulting in a white solid.
vii. **General procedure for MALDI-TOF of PLPs:** PLPs were dissolved in water (1 mg/mL) and added to a solution of α-cyano-4-hydroxycinnamic acid (50% acetonitrile, 0.1% TFA in DI water, saturated) in a ratio of 1:10. 1 µL of the sample-matrix solution was drop cast and allowed to dry prior to mass analysis. Samples were then analyzed using a Bruker rapileX Tissuetyper in negative and positive ion mode (1000-5000 shots, 1 kHz, laser beam attenuation 60-40%).
viii. **Enzyme degradation assay via analytical HPLC:** Both compounds were dissolved at the same concentration with respect to peptide (100 μM). The final concentration of peptide to enzyme was 20,000:1. The enzyme that was chosen was trypsin as it will selectively cleave the peptide at the N-terminus of the arginine. Both PLP and peptide were dissolved (TRIS buffered water, 10 mL), heated to 40°C, and added enzyme (acetic acid buffered water, 2.5 μL, 500 ug/mL). Aliquots of the reaction were taken across a week. Following deactivation of the enzyme using trifluoracetic acid, samples were run using analytical HPLC.
ix. **Human Corneal Epithelial Cell Culture:** Human corneal epithelial cells (hTCEpi) were incubated at 37˚C at 5% CO_2_ using Keratinocyte SFM medium supplemented with bovine pituitary extract (BPE, 25 mg), epidermal growth factor (EGF 1-53, 2.5 μg), antibiotics (Penicillin-Streptomycin, Life Technologies, 15140122), and calcium chloride (0.2 M, sterile filtered). Cell cultures were subculturing in flasks every 3-5 days when cells became 80% confluent using with trypsin-EDTA (0.05% Trypsin, 0.02% EDTA, PCS-999-003) and neutralized with 5% FBS (heat inactivated, Corning, 35011CV) in PBS ([-] calcium, magnesium, 45000-434).
x. **Cellular uptake via confocal microscopy:** For treatment and imaging, cells were seeded in 4-chamber, 35 mm round glass-bottom dishes at a density of 15,000 cells per well and incubated for 24 hours in a 5% CO_2_ atmosphere at 37°C before polymer treatment. PLP used for treatments was dissolved in D.I. Water and added into dish wells at the indicated concentrations (final concentration of PLP treatments 5 µM, 1 µM, and 0.5 µM), followed by 24-hour incubation; control wells received only D.I. water. Before confocal microscopy imaging, cells were washed with media to remove any PLP not taken up by the cells, ensuring that all fluorescence observed in the confocal is from PLP inside the cells. After washing, the cells were stained with blue Hoechst 33342, adding one drop per well. Cells were imaged in sets of triplicate (1 triplicate per well). Images were done using a Leica SP8 with excitation lasers at 405nm for the blue channel (blue Hoechst 33342), with a detection channel for the blue laser centered at 460 nm (+/- 10nm width range) and red laser emitting at 633nm and the detection centered at 660nm (+/- 10nm), for the Cye5.5. Bright-field images were illuminated using the 405nm blue laser diode. The Lens used was a 63x/1.40 HC PLAPO Oil CS2. Scans were performed at a 200 Hz speed and a 2048 x 2048-pixel resolution.
xi. **Nitrogen mustard (NM) injury model:** Six week old wild-type mice (C57BL/6) females purchased from Charles River) were anesthetized using ketamine (100 mg/kg) and xylazine (10 mg/kg). Mouse corneas were exposed to 2 mm sterile filter paper discs soaked in 0.5% solution of mechlorethamine hydrochloride (nitrogen mustard; Sigma) for one minute. Corneas were clinically examined using a slit lamp to determine corneal clarity. Corneal haze score was determined as previously described. Corneas of live mice were examined by a Heidelberg Engineering Optical Coherence Tomography (OCT) system, which is equipped with an anterior segment module. A single image was taken at the central cornea for central corneal thickness measurement. All procedures received approval from the Institutional Animal Care and Use Committee (IACUC) at Northwestern University and are in adherence to the ARVO Statement for the Use of Animals in Ophthalmic and Vision Research.
xii. **Treatments *in* *vivo*:** Twenty-four hours post-NM injury, mice were either injected intraperitoneally or topically applied with NLE Ang (1-7) PLP, NLE Ang (1-7) peptide, or saline (vehicle control) daily. Uninjured mice were used as control.
xiii. **Corneal penetration *in* *vivo*:** Twenty-four hours post-NM injury, mice were topically applied with fluorescently-labeled NLE Ang (1-7) PLP-Cy5. Mice were harvested at 1 hr., 4 hr., 8 hr., and 24 hr. Eyes were frozen using optical cutting temperature compound, and sectioned. These slides were then DAPI nuclear stained and imaged using fluorescence microscopy.
xiv. **Clinical Imaging:** To evaluate clinically for corneal clarity, a slit lamp examination was performed to visualize the corneal haze. Corneal haze was scored as follows 0, no corneal haze; 1, iris detail visible; 2, Pupillary margin visible, iris detail obscured; 3, pupillary margin not visible; 4, cornea totally opaque. Epithelial defects were evaluated by applying 0.5% fluorescein in PBS and imaging under a light microscope with blue light. Optical Coherence Tomography (OCT) imaging was done on day 7 and day 13 in delayed phase treatment to evaluate the cornea thickness.
xv. **Reverse transcription-quantitative polymerase chain reaction (RT-qPCR):** Mouse corneas were dissected out as described previously(7). Total RNA was isolated using a miRNeasy mini kit (Qiagen, Valencia, CA, USA). cDNA was synthesized using SuperScript™ III First-Strand Synthesis SuperMix for qRT-PCR (ThermoFisher Scientific). Real-time qPCR was performed using the Roche FastStart Essential DNA Green Master (Roche, USA) in a Bio-Rad real-time PCR System as previously described (8). The mice primer pairs for RT‐qPCR were purchased from IDT (Coralville, IA, USA).
xvi. **Whole mount staining:** Mouse corneas with limbus were dissected under stereoscope (7) and fixed in a fixative solution (1m of 10xPBS, 270µl of 37% formaldehyde, 10 µl of 2mM MgCl2, 200µl of 250mM EGTA pH 8.0, Water 8.25 mL) CD31 antibody (BD Biosciences), which detects vessels, and MUC5AC antibody (Thermo Scientific), which detects goblet cells, were used for staining as previously described (9). The corneas were imaged using Nikon ECLIPSE Ti2 Fluorescence Microscope as well as Nikon SoRa superresolution microscope system. The vascularized area and the area occupied by goblet cells were measured by ImageJ. Vascularization was determined by vascularized area / the total corneal area. Conjunctivalization was determined by the area occupied by goblet cells divided by the total corneal area.
xvii. **Histology:** Whole mouse eyes were collected and immersed in 10% neutral buffered formalin for 24 hours. Samples were subsequently processed for paraffin embedding, sectioned, and subjected to hematoxylin and eosin (H&E) staining. Sections were first deparaffinized and rehydrated, then stained with Mayer’s hematoxylin for 30 seconds and rinsed under running tap water for 2 minutes. Differentiation was performed by briefly dipping slides twice in 1% acid alcohol, followed by an additional 2-minute wash in tap water. Slides were then counterstained with eosin (1 mL eosin supplemented with 2 µL glacial acetic acid) for 2–3 minutes. After several rinses in tap water, sections were dehydrated and coverslipped. Microscopic evaluation was carried out using a Zeiss Axioplan 2 light microscope. Quantitative analysis of corneal epithelial thickness and stromal inflammatory cell infiltration was performed using ImageJ software.
xviii. **Biodistribution via ICPMS:** The mice were injured 24 hours prior to treatment. Mice were then dosed topically with 0.5 mg/kg in saline. At time points of 48 hr, 24 hr, 8 hr, and 1 hr, four mice were anesthetized and exsanguinated. Whole blood and livers were collected in preweighed Falcon tubes. The tubes were weighed again to determine total mass of tissue collected in each tube. To each tube of all samples except liver, 300 μl of conc nitric acid and 30% hydrogen peroxide were added. Liver samples had 500 μl of conc nitric acid and 30% hydrogen peroxide added. All samples were digested in a 65°C water bath for 24 hours until the solutions were transparent. The tubes were diluted to 5 mL of total volume (10 mL for liver) with Millipore water and weighed for a final time. After this point, any excess digested tissue that precipitated was removed by centrifugation and the supernatant saved. Each sample was run on high-resolution ICPMS, the Thermo iCAP Q ICPMS, which, after calibration with known Gd concentrations, resulted in exact Gd concentration in parts per billion (ppb) of each sample. The limit of detection is 0.04 ppb. The ppb of Gd were directly related to the calibration curve of known Gd PLP concentrations and was used to calculate the amount of PLP present in each sample.
xix. **Statistical Analysis:** Data was analyzed using GraphPad Prism (version 10). The Mann–Whitney U test and one-way ANOVA were performed to determine statistical significance. The differences were considered significant for P values of <0.05.

**List of Figures and Tables**

**Fig S1.** Chemical structure, ESI and analytical HPLC trace of **NLE Ang (1-7) peptide**

**Fig S2.** Chemical structure, ESI and analytical HPLC trace of **M1**

**Fig S3.** Polymerization kinetics of PLP

**Fig S4.** Characterization of PLP using SEC, DLS, and MALDI

**Fig S5.** HPLC traces of peptide and PLP enzyme degradation

**Fig S6.** Calibration curve and cleaved peptide (%) over time

**Fig S7.** Cellular uptake of PLP in human corneal epithelial cells

**Fig S8.** Fluorescence imaging of eyes treated with Cy5-PLP

**Fig S9.** RT-qPCR of 3 day IP or topical treatment compared to topical steroid

**Supporting Table 1.** Calculated and observed ESI

**Supporting Table 2.** Primers for qPCR

**Supporting Table 3.** ICP-MS of Blood and Liver after PLP-Gd administration

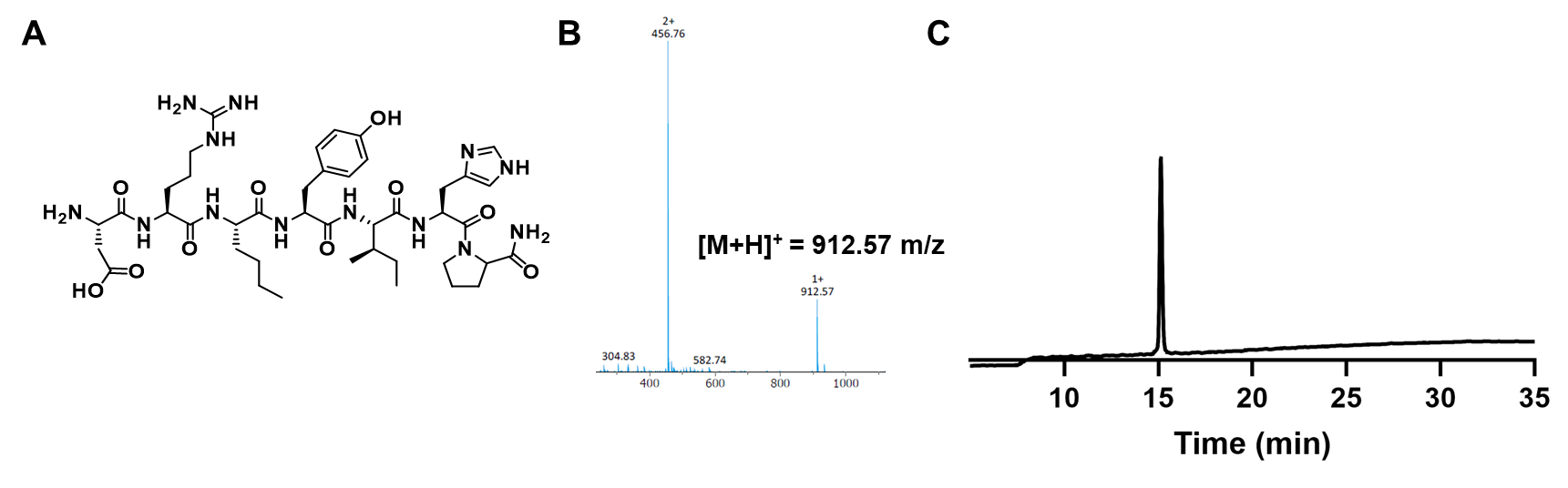

**Fig S1.** (A) Chemical structure, (B) ESI-MS and (C) analytical HPLC trace of

**NLE Ang (1-7) peptide.**

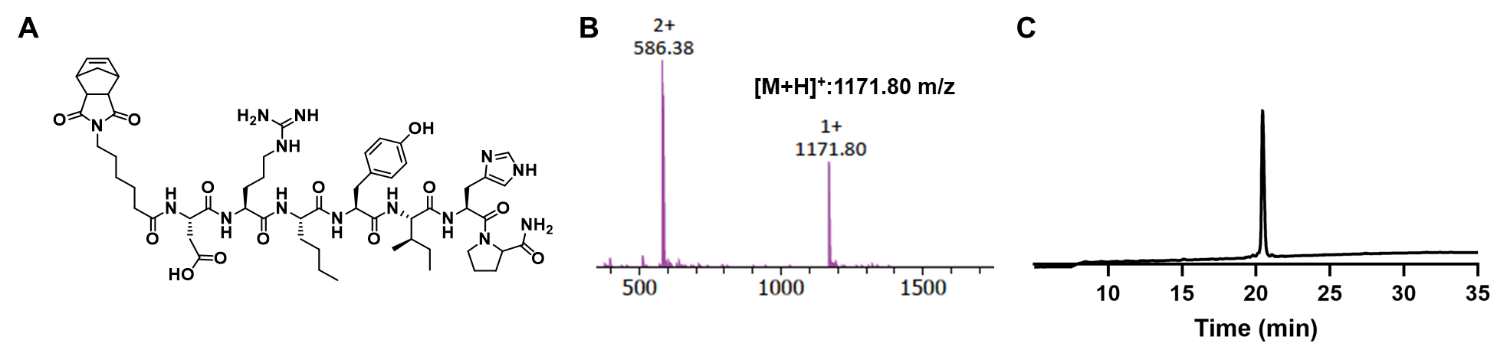

**Fig S2.** (A) Chemical structure, (B) ESI-MS and (C) analytical HPLC trace of **M1.**

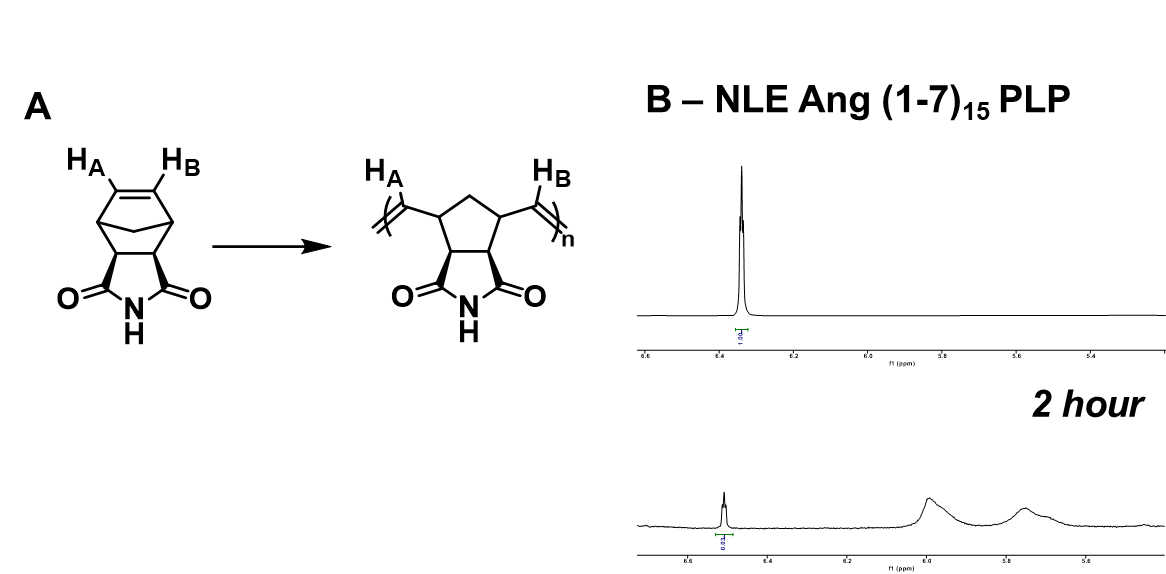

**Fig S3.** (A) Polymerization kinetics tracked via nuclear magnetic resonance spectroscopy (NMR). Kinetic experiments were performed by tracking the (A) disappearance of the monomer singlet peak (H_A_, H_B_, ~6.5ppm) and the appearance of the polymer multiplet peak (H_C_, H_D_, ~5.6ppm). (B) NLE Ang (1-7) PLP homopolymers with a degree of polymerization of 15 were polymerized over two hours (~97% monomer consumed).

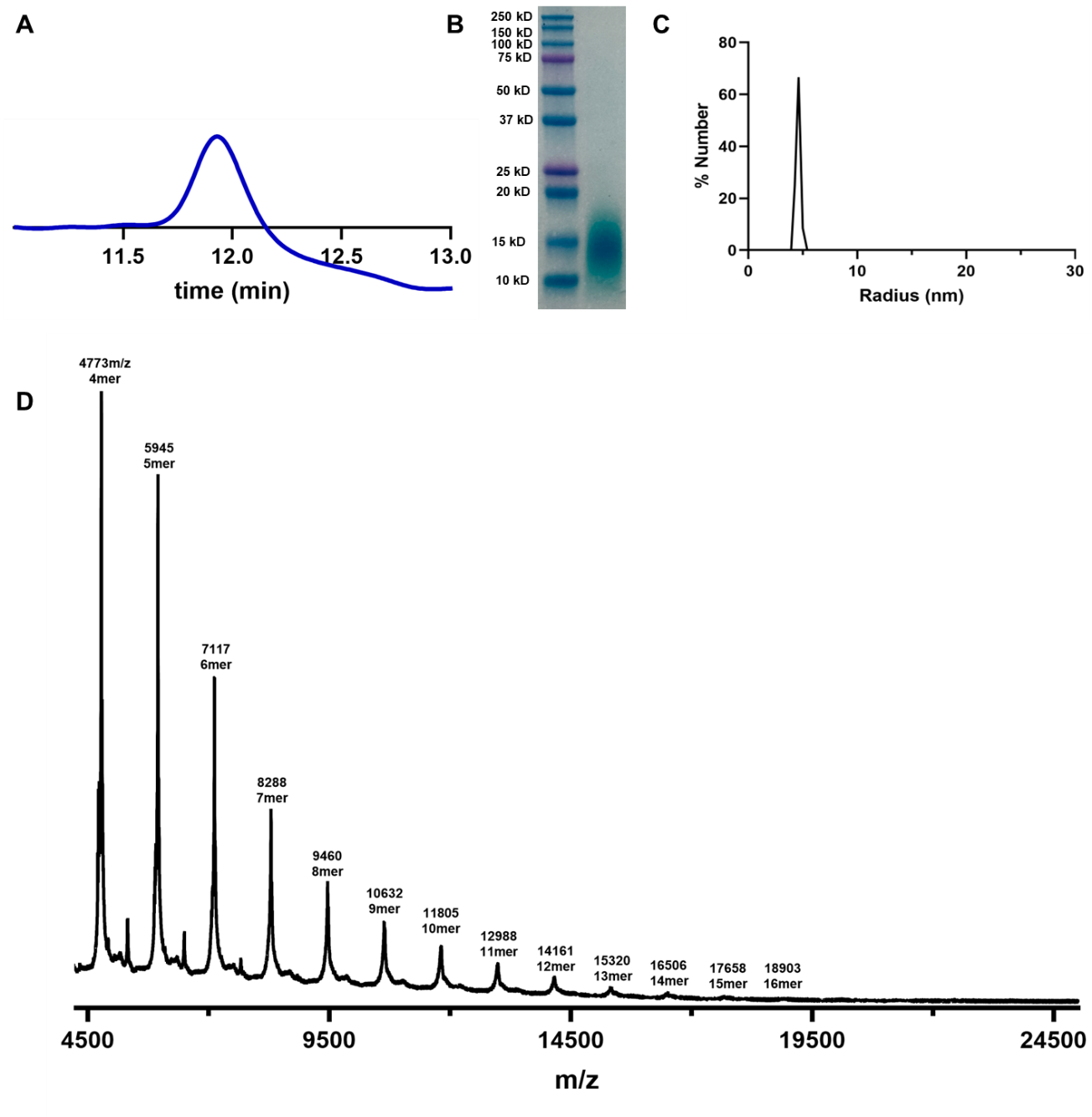

**Fig S4.** Characterization of NLE Ang (1-7)_15_ PLP. (A) The M_N_ via SEC-MALS is 16,000 Da and a dispersity (Đ) of 1.6. (B) SDS-PAGE shows molecular weight of 15 kDa. (C) DLS shows a hydrodynamic radius of 4 nm (1 mg/mL in saline). (D) PLPs are dropcast with 1:10 ratio of PLP to CHCA. MALDI spectrum has peaks spacing of ~1192 g/mol correlating to the MW of the monomer.

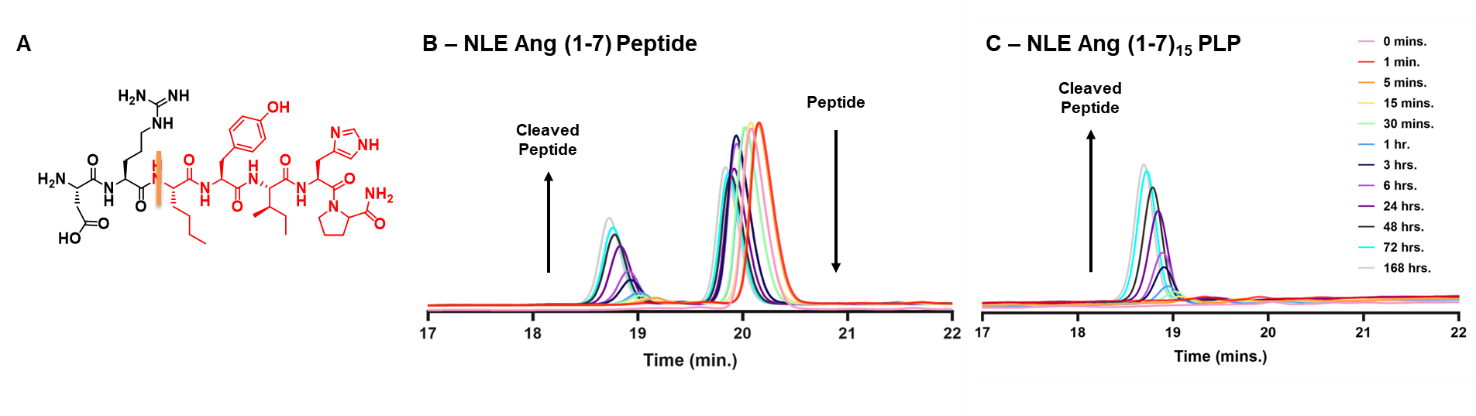

**Fig S5.** (A) Trypsin was added to the samples for selective cleavage at the C-terminus of the arginine residue. HPLC traces are shown of the cleaved peptide fragments of enzyme-treated (B) peptide and (C) PLP over a week.

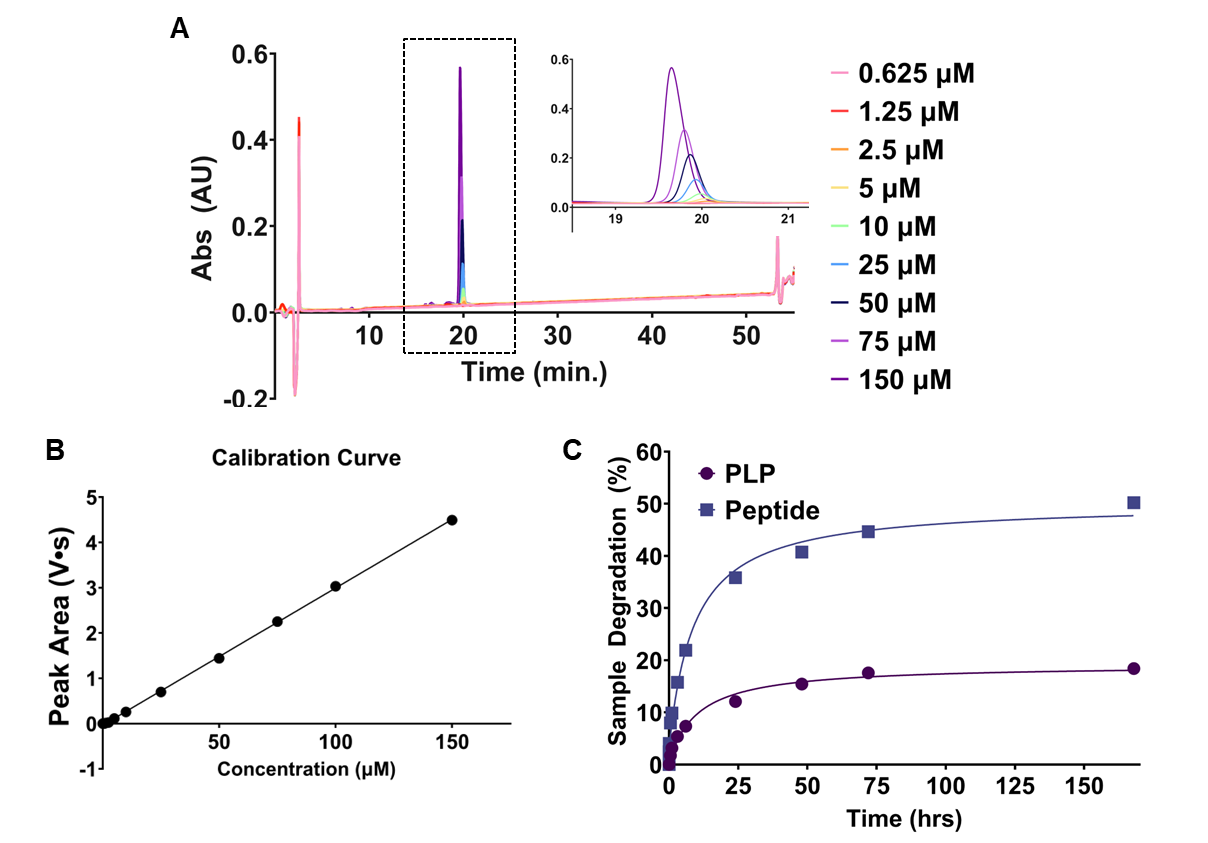

**Fig S6.** Quantification of trypsin degradation study (Fig S5). (A) To make a calibration curve to determine percent peptide cleaved, the peptide fragment (NleYIHP) was run on HPLC at concentrations ranging from 3.13-250 µM. (B) Area under the curve was plotted for each concentration. (C) Calibration curve was used to determine and plot cleaved peptide (%) over 72 hours for peptide and PLP.

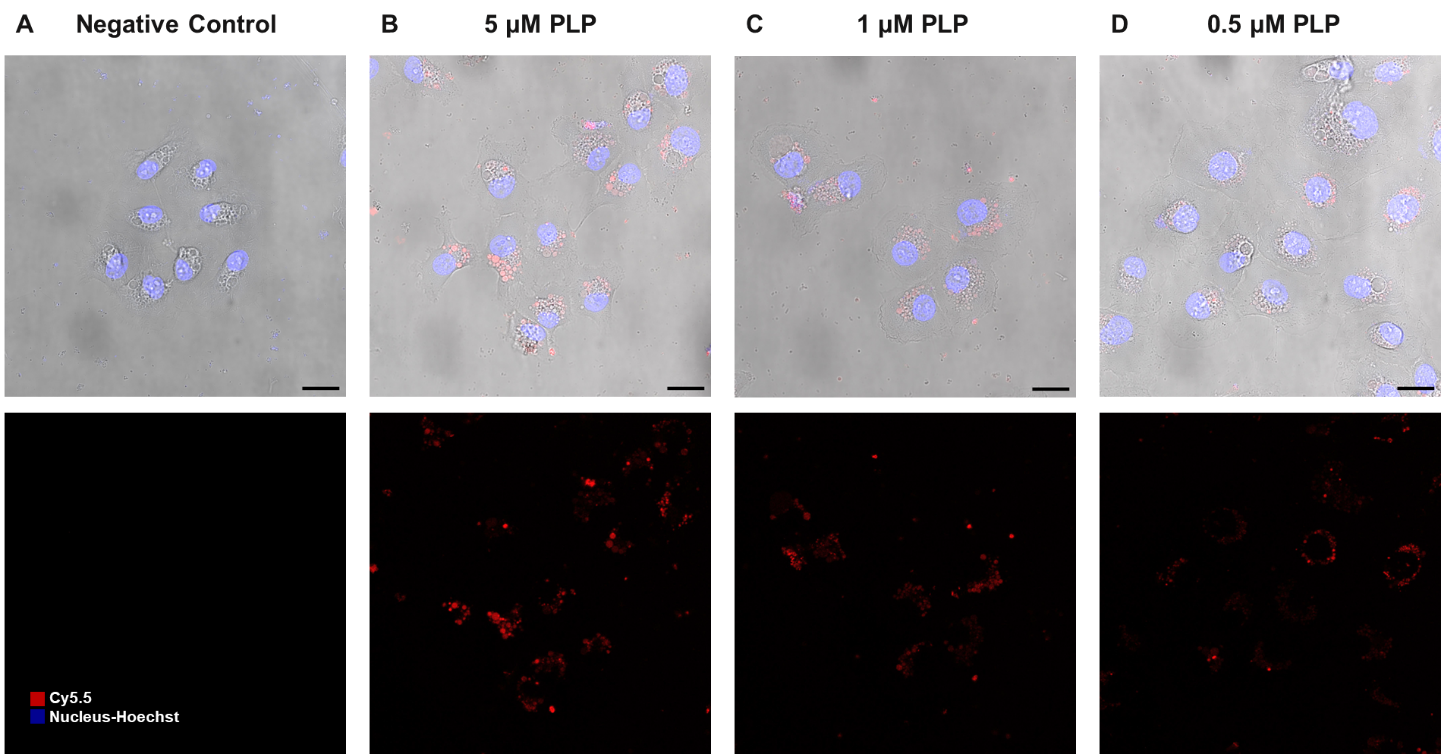

**Fig S7.** Cellular uptake of PLPs in corneal cells. PLPs were incubated in hTCEpi cells for 24 hrs. (A) Cells were treated with a nuclear stain (blue) and imaged. Cellular uptake of PLPs (red) was detected at (B) 5 µM, (C) 1 µM, and (D) 0.5 µM. Scale bar 40µm.

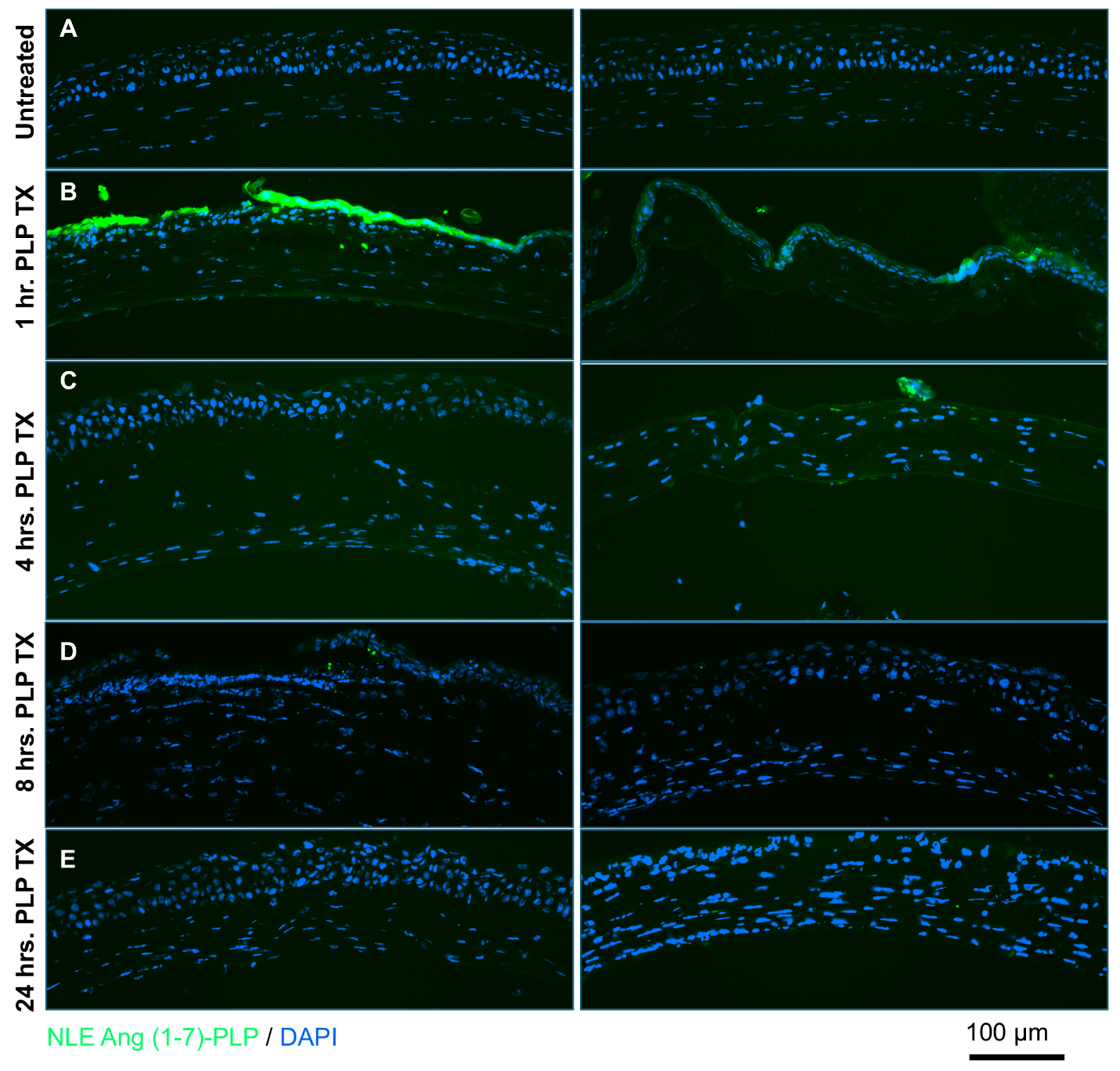

**Fig S8.** Mice (N=2) were injured and treated with fluorescently labeled PLPs (green) the following day. Slides sectioned and stained with DAPI (blue). Fluorescence was compared to (A) untreated controls. Mice were harvested at (B) 1 hr., (C) 4 hr., (D) 8 hr., and (E) 24 hr. The only fluorescence detected that we attribute to the PLP is at hour 1.

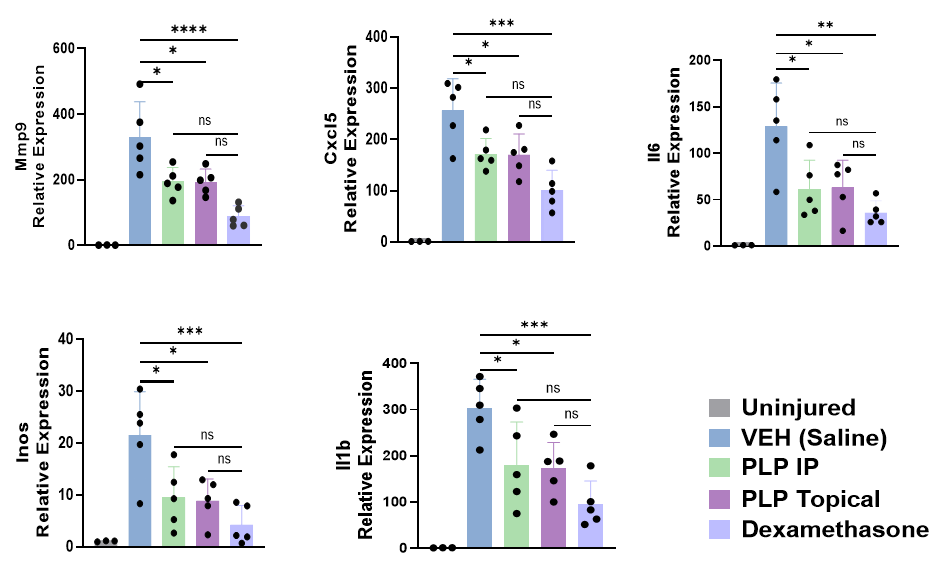

**Fig S9.** RT-qPCR revealed no significant difference in reduction of inflammatory gene expression between dexamethasone and PLP treated groups in acute phase of NM injury, indicating that PLP treatment achieved a comparable anti-inflammatory effect (n=5). * P ≤ 0.05, **:P ≤ 0.01, ***P ≤ 0.001, **** P ≤ 0.0001.

**Supplemental Table S1**. Calculated and observed mass via ESI for

NLE Ang (1-7) peptide and **M1**

| **Abbreviation** | **Monomer** | **Calculated Mass** | | **Obtained Mass (ESI)** |
| --- | --- | --- | --- | --- |
| NLE Ang (1-7) | DR(Nle)YIHP | | 911.50 | [M+H]^+^: 912.57 |
| M1 | NorAha^1^-DR(Nle)YIHP | | 1170.62 | [M+H]^+^: 1171.80 |

^1^Abbreviation for *N*-(hexanoic acid)-exo-5-norbornene-2,3-dicarboximide

**Supplemental Table S2.** Primers for qPCR

| **Gene** | **Direction** | **Primer Sequence (5'-3')** |
| --- | --- | --- |
| MMP9 | Forward | CCAGTACCAAGACAAAGCCTAT |
|  | Reverse | TCCACCTTGTTCACCTCATTT |
| iNOS | Forward | TGTGGCTACCACATTGAAGAA |
|  | Reverse | CATGATAACGTTTCTGGCTCTTG |
| IL-6 | Forward | GTATGAACAACGATGATGCACTTG |
|  | Reverse | GGCAAATTTCCTGATTATATCCAGTTTG |
| CXCL5 | Forward | CTA CGG TGG AAG TCA TAG CTA AA |
|  | Reverse | GCA TTC CGC TTA GCT TTC TTT |
| OSM | Forward | TCA CTC TTG GAG CCC TAT ATC C |
|  | Reverse | AAG TGA GGC TTG CTC AGT TG |
| Vegf | Forward | ATGCAGACCAAAGAAAGACAGAAC |
|  | Reverse | GAGAGGTCTGGTTCCCGAAA |
| Pdgfb | Forward | GAG TGT GGG CAG GGT TAT TT |
|  | Reverse | GAA TCA GGC ATC GAG ACA GAC |
| IL1b | Forward | CTGCAGCTGGAGAGTGTG |
|  | Reverse | CTCCACTTTGCTCTTGACTTCTA |

**Supplemental Table S3.** ICP-MS detection of PLP-Gd in blood and liver after eye drop administration across 48 hours.

| **Sample** | **Timepoint (hr.)** | **Tissue (g)** | **Gd in Tissue (ppb)** |
| --- | --- | --- | --- |
| 1 | 48 | Blood | 0.015 |
| 2 | 48 | Blood | 0.045 |
| 3 | 48 | Blood | 0.007 |
| 4 | 48 | Blood | 0.049 |
| 5 | 24 | Blood | 0.011 |
| 6 | 24 | Blood | 0.008 |
| 7 | 24 | Blood | 0.057 |
| 8 | 24 | Blood | 0.046 |
| 9 | 8 | Blood | 0.031 |
| 10 | 8 | Blood | 0.022 |
| 11 | 8 | Blood | 0.037 |
| 12 | 8 | Blood | 0.051 |
| 13 | 1 | Blood | 0.008 |
| 14 | 1 | Blood | 0.013 |
| 15 | 1 | Blood | 0.024 |
| 16 | 1 | Blood | 0.029 |
| 17 | Uninjured - NTC | Blood | 0.031 |
| 18 | Uninjured - NTC | Blood | 0.021 |
| 19 | Uninjured - NTC | Blood | 0.086 |
| 20 | Uninjured - NTC | Blood | 0.038 |
| 21 | Injured - NTC | Blood | 0.025 |
| 33 | 48 | Liver | 0.034 |
| 34 | 48 | Liver | 0.187 |
| 35 | 48 | Liver | 0.058 |
| 36 | 48 | Liver | 0.069 |
| 37 | 24 | Liver | 0.034 |
| 38 | 24 | Liver | 0.136 |
| 39 | 24 | Liver | 0.043 |
| 40 | 24 | Liver | 0.036 |
| 41 | 8 | Liver | 0.076 |
| 42 | 8 | Liver | 0.026 |
| 43 | 8 | Liver | 0.069 |
| 44 | 8 | Liver | 0.080 |
| 45 | 1 | Liver | 0.091 |
| 46 | 1 | Liver | 0.032 |
| 47 | 1 | Liver | 0.054 |
| 48 | 1 | Liver | 0.067 |
| 49 | Uninjured - NTC | Liver | 0.032 |
| 50 | Uninjured - NTC | Liver | 0.234 |
| 51 | Uninjured - NTC | Liver | 0.022 |
| 52 | Uninjured - NTC | Liver | 0.037 |
| 53 | Injured - NTC | Liver | 0.052 |

**References**

1.	M. S. Sanford, J. A. Love, R. H. Grubbs, A Versatile Precursor for the Synthesis of New Ruthenium Olefin Metathesis Catalysts. *Organometallics* **20**, 5314–5318 (2001).

2.	M. P. Thompson, *et al.*, Labelling polymers and micellar nanoparticles via initiation, propagation and termination with ROMP. *Polym. Chem.* **5**, 1954–1964 (2014).

3.	J. K. Kammeyer, A. P. Blum, L. Adamiak, M. E. Hahn, N. C. Gianneschi, Polymerization of protecting-group-free peptides via ROMP. *Polym. Chem.* **4**, 3929–3933 (2013).

4.	C. Battistella, *et al.*, Delivery of Immunotherapeutic Nanoparticles to Tumors via Enzyme-Directed Assembly. *Adv. Healthc. Mater.* **8**, 1901105 (2019).

5.	W. Choi, *et al.*, Thrombospondin-1 proteomimetic polymers exhibit anti-angiogenic activity in a neovascular age-related macular degeneration mouse model. *Sci. Adv.* **9**, eadi8534 (2023).

6.	W. Choi, *et al.*, Proteomimetic polymer blocks mitochondrial damage, rescues Huntington’s neurons, and slows onset of neuropathology in vivo. *Sci. Adv.* **10**, eado8307 (2024).

7.	E. Kayaalp Nalbant, *et al.*, A novel therapy to ameliorate nitrogen mustard-induced limbal stem cell deficiency using lipoprotein-like nanoparticles. *NPJ Regen. Med.* **10**, 14 (2025).

8.	M. Notara, A. Lentzsch, M. Coroneo, C. Cursiefen, The Role of Limbal Epithelial Stem Cells in Regulating Corneal (Lymph)angiogenic Privilege and the Micromilieu of the Limbal Niche following UV Exposure. *Stem Cells Int.* **2018**, 8620172 (2018).

9.	J. Wang, *et al.*, The ACE2‐deficient mouse: A model for a cytokine storm‐driven inflammation. *FASEB J.* **34**, 10505–10515 (2020).
